# Proteomics unveil a central role for peroxisomes in butyrate assimilation of the heterotrophic Chlorophyte alga *Polytomella* sp

**DOI:** 10.1101/2022.08.29.505702

**Authors:** J. Lacroux, A. Atteia, S. Brugière, Y. Couté, O. Vallon, J.-P. Steyer, R. van Lis

## Abstract

Volatile fatty acids found in effluents of the dark fermentation of biowastes can be used for mixotrophic growth of microalgae, improving productivity and reducing the cost of the feedstock. Microalgae can use the acetate in the effluents very well, but butyrate is poorly assimilated and can inhibit growth above 1 gC.L^-1^. The non-photosynthetic chlorophyte alga *Polytomella* sp. SAG 198.80 was found to be able to assimilate butyrate fast. To decipher the metabolic pathways implicated in butyrate assimilation, quantitative proteomics study was developed comparing *Polytomella* sp. cells grown on acetate and butyrate at 1 gC.L^-1^. After statistical analysis, a total of 1772 proteins were retained, of which 119 proteins were found to be overaccumulated on butyrate vs. only 46 on acetate, indicating that butyrate assimilation necessitates additional metabolic steps. The data show that butyrate assimilation occurs in the peroxisome via the β-oxidation pathway to produce acetyl-CoA and further tri/dicarboxylic acids in the glyoxylate cycle. Concomitantly, reactive oxygen species defense enzymes as well as the branched amino acid degradation pathway were strongly induced.Although no clear dedicated butyrate transport mechanism could be inferred, several membrane transporters induced on butyrate are identified as potential condidates. Metabolic responses correspond globally to the increased needs for central cofactors NAD, ATP and CoA, especially in the peroxisome and the cytosol.

## 1 Introduction

Mixotrophic growth combines the reduction of CO_2_ via photosynthesis with the oxidation of organic carbon and is found in a wide range of phototrophic microorganisms such as cyanobacteria and microalgae (Perez-Garcia and Bashan, 2015). This trophic mode is generally the most efficient in terms of biomass productivity (Zhan et al., 2017). Some photosynthetic microalgae also have heterotrophic capacities, i.e. they can grow in the absence of light on reduced carbon sources (Round, 1980). Although glucose can yield very high biomass productivities (Perez-Garcia et al., 2011), it is an expensive substrate that is not economical for many microalgal biorefinery applications with lower added value such as biofuels, green chemistry platform molecules, aquaculture and biofertilizer (Acién Fernández et al., 2019). It has been proposed to use organic acids produced by the fermentation of biowaste material as feedstock to lower the cost (Delrue et al., 2016; Turon et al., 2016; Chalima et al., 2017; Karnaouri et al., 2020). Dark fermentation (DF) of organic matter by microbial consortia is a sustainable n method for H_2_ production that compares favorably to other process in terms of CO_2_ footprint (Dincer and Acar, 2015; Moscoviz et al., 2018). In this context it is warranted to intensify research on downstream coupled processes such as mixotrophic cultivation of microalgae on DF effluents (DFE). Depending on the conditions, DFEs contain different ratios of volatile fatty acids (VFA) alongside some lactate and ethanol. In general, mostly acetate and butyrate are present, in a molar ratio of 0.66 on average (Turon et al., 2016; Moscoviz et al., 2018).

While microalgae that use glucose can usually also use acetate, many algae can grow on acetate but not on glucose, e.g. *Chlamydomonas reinhardtii* (Harris, 1989). Capacities for the use of other substrates such as lactate, ethanol and butyrate are less common and vary widely, even within the same species e.g. *Euglena gracilis var bacillaris* or *urophora* (Neilson and Lewin, 1974; Hosotani et al., 1988). By far the most abundant VFA in DFEs is butyrate, but it is toxic for many bacteria and is known to be poorly used by microalgae (Turon et al., 2015; Lacroux et al., 2020, 2021). A generalized toxicity effect of VFAs is observed as the extracellular pH approaches the pKa value because cells are permeable to the protonated form, resulting in deleterious effect on the cell metabolism and integrity (Lacroux et al., 2020). The toxicity threshold for butyrate (as butyric acid) is 5-fold lower than for acetic acid in some species, explaining why it is a poor carbon source. Algal growth on butyrate will thus require more care in adapting the pH and concentration to remain below the observed species-specific toxicity threshold. However, this will be inadequate to reach growth rates similar to those on acetate. It is therefore crucial that butyrate metabolism be studied in detail in order to remove metabolic bottlenecks in its assimilation.

Butyrate assimilation is well studied in microorganisms such as the sulfate reducer *Desulfosarcina cetonica* (Janssen and Schink, 1995), the non-sulfur purple bacterium *Rhodospirillum rubrum* (De Meur et al., 2020), yeasts i.e. *Candida ingens* (Garrison et al., 1985) or *Yarrowia lipolytica* (Llamas et al., 2020) and also in human colonocytes, for which bacteria-produced butyrate is the primary energy source (Roediger, 1982; Fleming et al., 1991). Colonocytes import butyrate via the monocarboxylate transporter (MCT), a 45-kDa plasma membrane protein with 12 transmembrane segments that symports H+ with the butyrate anion (Cuff et al., 2005). Butyrate is subsequently imported into the mitochondrial matrix where it undergoes β-oxidation to acetyl-CoA, which in turn enters the TCA cycle resulting in the production of NADH. The first step of β-oxidation is its activation into butyryl-CoA via an ATP-dependent butyryl-CoA synthetase, followed by the conversion into crotonyl-CoA by butyryl-CoA dehydrogenase, into hydroxy-isobutyryl-CoA by enoyl-CoA hydratase, then into acetoacetyl-CoA by hydroxybutyryl-CoA dehydrogenase and finally into acetyl-CoA via acetoacetyl-CoA thiolase (De Preter et al., 2012). A different assimilation metabolism was uncovered by a proteomic approach in the non-sulfur purple bacterium *R. rubrum* (De Meur et al., 2020), where acetyl-CoA is used to activate butyrate via butyryl-CoA:acetate CoA transferase under photoheterotrophic conditions. Homologs of most enzymes potentially involved in butyrate assimilation as found in human colonocytes have been identified in *C. reinhardtii*, the best studied algal species (Li-Beisson et al., 2019). However, nothing is known about the enzyme(s) that may be involved in formation of butyryl-CoA in algae, in particular their subcellular localization. The case of acetyl-CoA is better studied. In the model alga *C. reinhardtii*, acetyl-CoA is formed from acetate by acetyl-CoA synthase, purportedly in peroxisomes, and further utilized in the glyoxylate cycle to form products such as malate, citrate and succinate that can be further exported to other cell compartments (Lauersen et al., 2016). The import of butyrate into microalgal cells and the associated metabolic responses and their intracellular localization remain to be studied.

The heterotrophic chlorophyte *Polytomella* sp. SAG 198.80 is to our knowledge the alga for which the fastest butyrate assimilation has been described (Wise, 1955, 1959), and more recently by the authors of this work (Lacroux et al. 2022). The fact that *Polytomella* has lost photosynthetic activity allows focusing on the assimilation pathways, avoiding interactions with photosynthetic metabolism that complicate analysis (van Lis and Atteia, 2004; Johnson and Alric, 2013). In this study, a quantitative proteomics approach is used to decipher the metabolic pathways specifically involved in the assimilation of butyrate by *Polytomella*, based on the comparison to acetate as reference metabolism as this is the simplest entry of organic carbon into central carbon metabolism.

## 2 Material and methods

### 2.1 Strain and culture conditions

*Polytomella* sp. SAG 198.80 was obtained from the SAG culture collection (Goettingen, Germany). It was grown on synthetic media referred to as HAP (acetate) or HBP (butyrate), based on Tris-Acetate-Phosphate (TAP) medium used for the green alga *C. reinhardtii* (Harris, 1989), in which the Tris buffer is replaced by HEPES 0.1 M, at pH 7. Beijerincks solution (40X) was used at 25 mL.L^-1^ leading to an ammonium (NH_4_^+^) concentration of 7.5 mM, 0.6 mM of MgSO_4_ and 0.3 mM of CaCl_2_. To adjust nutrients to the Redfield C:N:P ratio of 106:16:1, corresponding to 83.3:12.6:0.8 mM for 1 g carbon per liter (1.0 g_C_.L^-1^), the proper amount of 1 M NH_4_Cl (5 mL) and 1 M K_2_HPO_4_ (0.8 mL) stock solutions were added. Hutner’s trace elements were used at 1 mL.L^-1^ (Hutner, 1972). As carbon source, acetate or butyrate were added as sodium salts at 1.0 g_C_.L^-1^ (41.7 mM acetate, 20.8 mM butyrate) and pH medium was adjusted to 7.0 (HCl) prior to sterilization at 121°C for 20 min. After cooling, 100 µL.L^-1^ was added of a stock of vitamin B1 (50 mM), biotin (1 mM) and cyanocobalamin (1 mM), sterilized over a 0.2 µm filter. Precultures were maintained on HAP medium containing 20 mM HEPES and were used to inoculate HAP and HBP media to initial optical density at 750 nm (OD_750_) = 0.05. The inoculum was prepared by collecting preculture cells in their exponential phase via centrifugation at 2500 g for 10 min, and resuspending them in Phosphate Buffered Saline to a final OD_750_ = 5. Cultures were done in 500mL Erlenmeyer flasks filled with 200 mL medium under dim light at 25°C and without agitation.

### 2.2 Biomass and VFA measurement

Biomass growth was followed by measuring optical density at 750 nm (OD750), using a Helios Epsilon spectrophotometer. OD750 was measured by placing 1 mL of liquid culture in a cuvette and comparing to distilled water. The sample was diluted when necessary so that OD750 < 0.6. Biomass production was expressed in g.L^-1^ dry weight (DW) deduced from OD750 determination. To calculate the biomass DW from OD750 values, a correlated factor was used, determined to be 1.0774 (R² = 0.977). This correlation factor was obtained from a calibration curve gathering 70 data points collected during separated experiments, at various growth stages (stationary phase excluded) and growth conditions (HAP or HBP medium). To determine biomass DW during these experiments, between 5 and 25 mL (depending on growth stage) biomass was centrifuged (3000 rpm, 10 min). Supernatant was discarded, and pellet rinsed with one volume of phosphate buffer saline. Biomass was centrifuged again, supernatant discarded and pellet resuspended in 10 mL distilled wated. The biomass was transferred in a pre-dried and pre-weighed aluminium crucible and dried overnight at 105°C.

For VFA measurements, samples from fresh cultures were immediately centrifuged, the supernatant filtered over 0.2 μm cut-off filters and frozen at -25°C until analyzed. The VFA concentrations were determined by gas chromatography. A 500 μL aliquot of supernatant was mixed with 500 μL of internal standard solution (ethyl-2-butyric acid, 1 g·L^-1^). The GC system consisted in a Perkin Clarus 580 model equipped with capillary column Elite-FFAP crossbond®carbowax® (15 m) maintained at 200 °C and with N_2_ as the gas vector (flow rate of 6 mL·min^-1^) with a flame ionization detector (FID) maintained at 280 °C (PerkinElmer, USA)

### 2.3 Analysis of total lipids and sugars

Samples from fresh cultures were immediately centrifuged, the supernatants were discarded and the pellets stored at -25°C until used. Prior to analysis, the pellet was thawed, resuspended in 100µl distilled water and added to 10 mL glass tubes for either lipid or sugar measurement. Total lipid concentrations in the algal samples were determined by the phosphovanillin method (Mishra et al., 2014). Phosphovanillin reagent was freshly prepared by first dissolving 0.6 g vanillin in 10 ml ethanol, and then adding 90 ml deionized water and 400 ml of H_3_PO_4_ (85%). The resulting reagent was stored in the dark. First, 2 mL H_2_SO_4_ (98%) were added in the tubes containing microalgae cells. The tubes were heated 10 min at 100° C and cooled on ice. The reaction was initiated by addition of 5 mL phosphovanillin reagent prior incubation for 15 min at 37° C. Tubes were periodically shaken by inversion. After cooling, optical density of suspensions was measured at 530 nm with an Aqualytic® spectrometer and compared to distilled water. Calibration curves were obtained using canola oil.

Total sugars were measured by the anthrone method (Yemm and Willis, 1954). Anthrone reagent was prepared by dissolving 200 mg of anthrone in 100 mL of H_2_SO_4_ (98%). Two mL of anthrone reagent were added in the tubes containing microalgae. Tubes were cooled down on ice and then incubated at 100° C for 10 min. After cooling, absorbance was measured at 625 nm with an Aqualytic® spectrometer and compared to distilled water. Calibration curves were obtained using glucose solution.

### 2.4 Calculation of specific rates and product yield in growing cultures

The biomass productivity P_x_ (g_dw_.L^-1^.d^-1^) and the specific growth rate *μ*_*x*_ (d^-1^) were calculated according to equations (1) and (2):

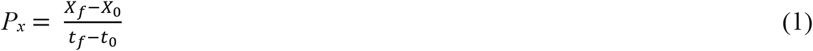

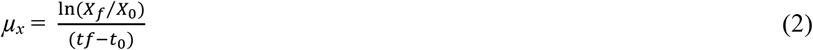

where *X*_*f*_ and *X*_*0*_ are the biomass concentrations at t=final (h) and at t=0h. The biomass yield Y (g_dw_.g_sub_^-1^) was estimated according to equation (3):

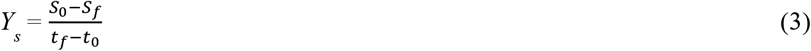

Where *S*_*f*_ and *S*_*0*_ are substrate concentrations (g.L^-1^) at t=final (h) and at t=0h. Statistical analysis was performed using GraphPad Prism V 8.0.2.

### 2.5 Mass spectrometry-based proteomic analyses

The total proteome of *Polytomella* sp. grown either on acetate or butyrate (three biological replicates per condition) were stacked in the top of a 12% SDS-PAGE resolving gel before in-gel digestion using modified trypsin (sequencing grade, Promega), as previously described (Casabona et al., 2013). The resulting peptides were analyzed by online nanoliquid chromatography coupled to MS/MS (Ultimate 3000 RSLCnano and Q-Exactive HF, Thermo Fisher Scientific) using a 80 min gradient. For this purpose, the peptides were sampled on a precolumn (300 μm x 5 mm PepMap C18, Thermo Scientific) and separated in a 75 μm x 250 mm C18 column (Reprosil-Pur 120 C18-AQ, 1.9 μm, Dr. Maisch). The MS and MS/MS data were acquired by Xcalibur (Thermo Fisher Scientific).

Peptides and proteins were identified by Mascot (version 2.6.0, Matrix Science) through concomitant searches against a homemade *Polytomella* sp. database, a homemade classical database containing the sequences of classical contaminant proteins found in proteomic analyses (human keratins, trypsin, etc.), and the corresponding reversed databases. Trypsin/P was chosen as the enzyme and two missed cleavages were allowed. Precursor and fragment mass error tolerances were set at respectively at 10 ppm and 25 mmu. Peptide modifications allowed during the search were: Carbamidomethyl (C, fixed), Acetyl (Protein N-term, variable) and Oxidation (M, variable). The Proline software (Bouyssié et al., 2020) was used for the compilation, grouping, and filtering of the results (conservation of rank 1 peptides, peptide length ≥ 7 amino acids, peptide-spectrum-match score ≥ 25, allowing to reach a false discovery rate of peptide-spectrum-match identifications < 1% as calculated on peptide-spectrum-match scores by employing the reverse database strategy, and minimum of one specific peptide per identified protein group). Proline was then used to perform MS1 quantification of the identified protein groups based on specific peptides.

Statistical analysis was performed using the ProStaR software (Wieczorek et al., 2017). The mass spectrometry proteomics data have been deposited to the ProteomeXchange Consortium via the PRIDE (Perez-Riverol et al., 2022) partner repository with the dataset identifier PXD035155. Proteins identified in the contaminant database, and proteins detected in less than three replicates of one condition were removed. After log2 transformation, abundance values were normalized by the vsn method before missing value imputation (slsa algorithm for partially observed values in the condition and DetQuantile algorithm for totally absent values in the condition). Statistical testing was conducted with limma, whereby differentially expressed (DE) proteins were sorted out using a log2(fold change) cut-off of 0.6 and a p-value cut-off of 0.004, allowing to reach a false discovery rate < 1% according to the Benjamini-Hochberg method.

Intensity-based absolute quantification (iBAQ, Schwanhäusser et al., 2011) values were calculated from MS intensities of specific peptides. For each sample, the iBAQ value of each protein was normalized by the summed iBAQ value of all proteins, before summing the values of the three replicates to generate the final iBAQ value of each condition.

### 2.6 Bioinformatic analyses

The assembly and structural annotation of the genome sequence of *Polytomella* sp. was described previously (van Lis et al., 2021). The predicted proteins (genome to be released at an later date) were used for the construction of the database for proteomics analysis. The functional annotation of the proteins that were identified by proteomics (1772 sequences) and further attribution of metabolic categories was done using Mercator4 V3.0 (https://plabipd.de/portal/mercator4), and used to project the fold change data onto a metabolic overview SVG image file, using the MapMan program (https://plabipd.de/portal/mapman) (Schwacke et al., 2019) used as stand-alone desktop application and Inkscape (https://inkscape.org/release/inkscape-1.2.1/) to modify the image file.

Final metabolic pathway reconstructions were based on metabolic maps made in KEGG mapper using KEGG and EC codes from BlastKOALA and KofamKOALA (HMM) homology searches at KEGG (https://www.genome.jp/kegg/) and by using EggNOG (Huerta-Cepas et al., 2019). These programs were also used to confirm and adjust annotations from Mercator where necessary, as well as manual BLAST searches using the UniProt database (https://www.uniprot.org) and the *C. reinhardtii* genome at Phytozome 13 (https://phytozome-next.jgi.doe.gov). OmicsBox 1.3.11 (https://www.biobam.com/omicsbox) genome blast was used with the genomes of *Chlamydomonas reinhardtii* (Uniprot UP000006906) and *Chlorella sorokiniana* (Uniprot UP000239899) to obtain equivalent enzyme identities in these two model microalgae. For subcellular localization predictions, DeepLoc (Almagro Armenteros et al., 2017) and PredAlgo (Tardif et al., 2012) were used with the limitation that the latter does not predict peroxisomal targeting. Therefore further searches for potential peroxisome targeting signals in *Polytomella* protein sequences were done with the PTS1 predictor (https://mendel.imp.ac.at/pts1/) (Neuberger et al., 2003) or manually based on (Gonzalez et al., 2011).

## 3 Results & Discussion

### 3.1 *Polytomella* sp. is a suitable model for the study of butyrate metabolism

At the basis of this work is the choice of *Polytomella* sp. as model algal species for the study of butyrate metabolism. The genus *Polytomella* belongs to the Reinhardtinia clade of Volvocine algae (Craig et al., 2021) and has diverged from a *Chlamydomonas*-like ancestor after having lost photosynthesis along with the chloroplast genome (Smith and Lee, 2014). *Chlamydomonas reinhardtii* only grows efficiently on acetate (Harris, 2001), so the highly versatile heterotrophic metabolism allowing *Polytomella* to grow on a multitude of alcohols and organic acids including butyrate (Wise, 1955, 1959; Round, 1980; de la Cruz and Gittleson, 1981) may have been partly acquired after the divergence. The availability of an annotated genome sequence of *Polytomella* sp. (van Lis et al., 2021) allowed a global proteomics approach to decipher its metabolic pathways involved in the assimilation of butyrate. Cells grown on either acetate or butyrate, at a fixed C concentration of 1.0 g_C_.L^-1^ (Fig. 1) showed similar growth rates of 2.37 d^-1^ ± 0.07 and 2.23 d^-1^ ± 0.08, respectively (three biological triplicates). Both carbon sources were completely consumed before 1.5 days, leading to a maximum biomass yield of 1.07 g_X_.L^-1^ (x=dry weight biomass) in both conditions (Fig 1C). After organic carbon exhaustion, biomass declined due to cell death in both conditions and cysts formation was observed (not shown). Biomass growth was accompanied by sugar accumulation, with about 15% less sugar on butyrate than on acetate (Fig 1D). In the related species *P. agilis*, the accumulated sugar was found to be essentially starch (Sheeler et al., 1968). Lipid accumulation is low in both cells, with a maximum of 73.3 ± 3.6 mg.L^-1^ (biomass content ∼5%) on acetate and 55.6 ± 3.1 mg.L^-1^ on butyrate.

**Figure 1.**
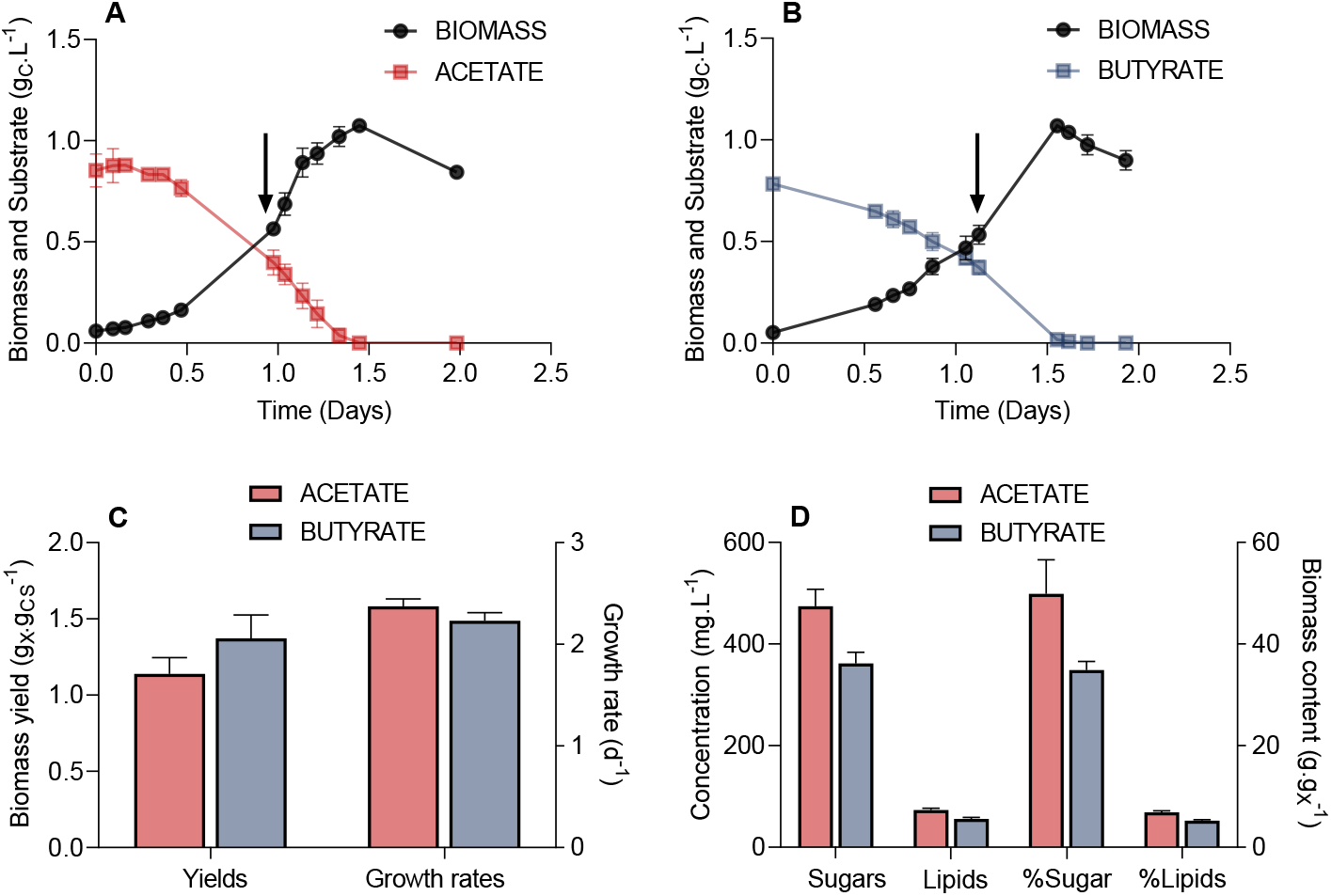
Parameters of the growth of P*olytomella* sp. on acetate and butyrate. A,B) Growth curves (g_X_.L^-1^) and substrate consumption (g_CS_.L^-1^) of *Polytomella* sp. in presence of 1 g_CS_.L^-1^ acetate or butyrate, C) biomass yields (g_X_.g_CS_^-1^) and growth rates (d^-1^) derived from these growth curves and D) the concentrations of sugars and lipids are plotted on the left (g.L^-1^) while their proportions to the total biomass are plotted on the right (g.g_X_^-1^) for both conditions. Arrows indicate when biomass for further proteomics analysis was sampled. X, dry weight; CS, dissolved organic carbon. Error bars correspond to standard deviations based on 3 biological replicates.

For several *Chlorella* strains it was reported that butyrate assimilation, when it occurred, was slower than acetate assimilation (Liu et al., 2013; Fei et al., 2015). Heterotrophic (dark) growth rates of *Chlorella sorokiniana* are 2.23 d^-1^ on acetate and 0.16 d^-1^ on butyrate (Turon et al., 2015). The ability of *Polytomella* sp. to grow on butyrate with a growth rate of 2.2 d^-1^, similar to that on acetate (Fig 1C), is thus remarkable.

Green microalgae tend to accumulate proteins during exponential growth and only start synthetizing storage compounds (starch, lipids) when growth conditions become suboptimal, e.g. when a micro- or macro-element such as nitrogen is limiting (Dincer and Acar, 2015). Since this fact usually implies that high biomass productivity is accompanied by low starch or lipid accumulation, continuous cultivation is less relevant (Adams et al., 2013). Accumulation of high levels of starch during the exponential growth phase is thus another distinctive and interesting trait of *Polytomella* sp..

### 3.2 Comparing global proteomes

Cultures of *Polytomella* sp. growing on acetate and butyrate were sampled during the exponential phase (arrows in Fig. 1). Total cell proteins were first analyzed on SDS-PAGE stained with Coomassie Blue G250. From the comparison of the protein profiles (Fig. 2A) it is inferred that butyrate elicits major changes in protein levels, and a clear upregulation of 3 proteins of 28, 38 and 60 kDa can be observed (indicated by asterisks, Fig 2A). To study more in detail the proteomic responses to butyrate, total proteins from acetate and butyrate grown cells were subjected to mass spectrometry (MS)-based label-free quantitative proteomic analysis. A volcano plot depicts the acetate vs butyrate fold change (FC, a direct measure of the relative protein abundance between both conditions) and the associated statistical significance (limma p-value) for each protein (Fig. 2B). The log2(FC) values varied between 4 and - 10, corresponding to an FC of 16 and 0.001, where the latter value indicates a 1000-fold upregulation on butyrate. A total of 1772 proteins were retained after statistical analysis, of which 119 were found to be significantly more abundant on butyrate compared to only 46 on acetate, and usually with a lower FC. This shows that growth on butyrate is accompanied by overaccumulation of a larger set of proteins than on acetate, which may indicate that butyrate needs more metabolic steps to enter central carbon metabolism than acetate.

**Figure 2.**
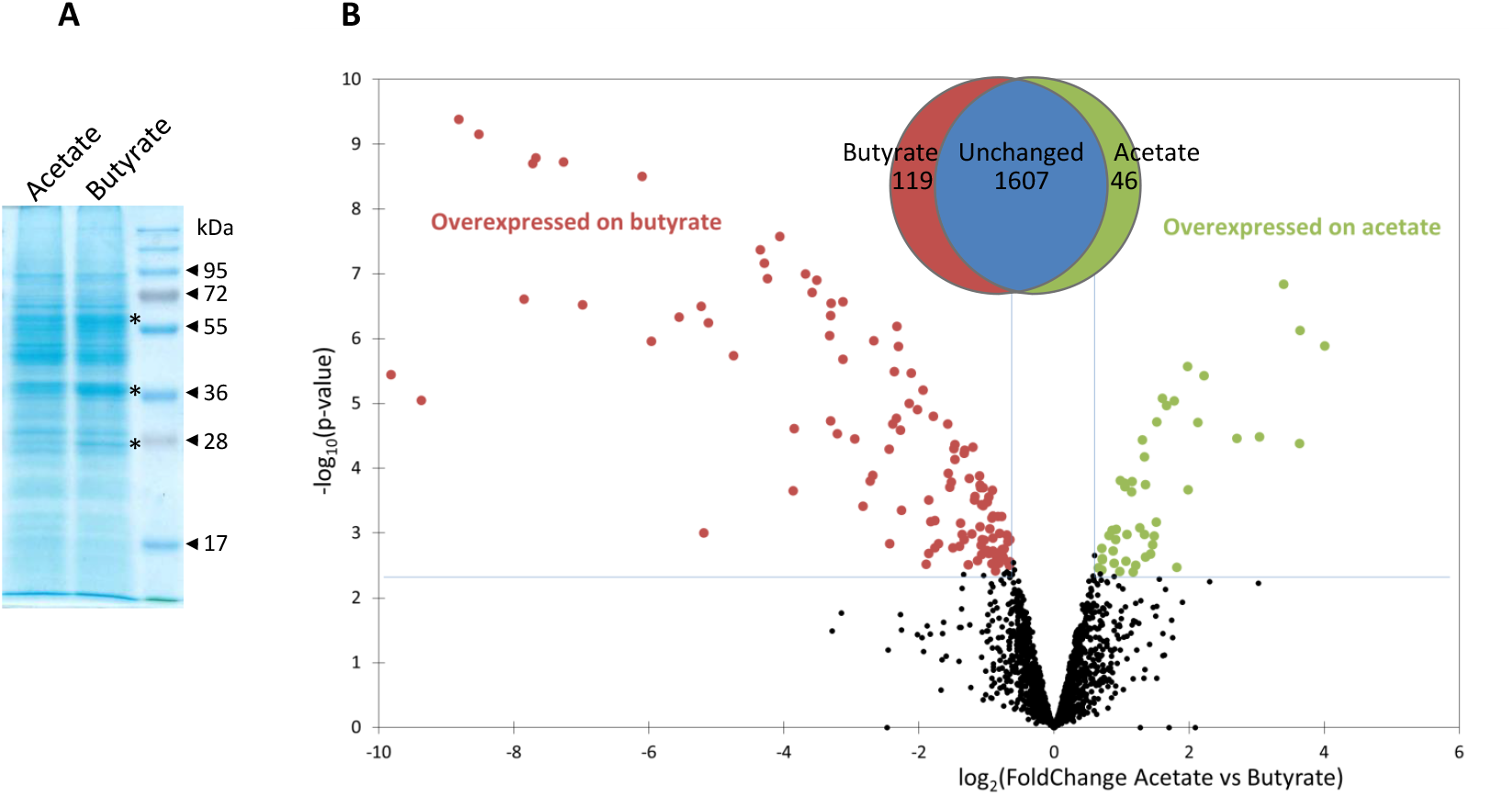
Comparison of global proteomes of *Polytomella* sp. grown on acetate and butyrate. A) Total proteins (30 µg) of *Polytomella* sp. growing exponentially on acetate or butyrate, resolved in a 12% SDS-polyacrylamide gel stained with Coomassie Blue G250. B) Volcano plot displaying the differential abundance of proteins of *Polytomella* sp. grown on acetate or butyrate analysed by MS-based quantitative proteomics. The volcano plot represents the -log10(limma p-value, cut off 0.004) on y-axis plotted against the log2(FoldChange acetate/butyrate) on the x-axis. Green and red dots represent proteins found more abundant in *Polytomella* sp. grown respectively on acetate or butyrate (Benjamini-Hochberg FDR < 1%). The Venn diagram indicates that of the total of 1772 proteins detected for acetate and butyrate combined, 48 were significantly induced on acetate and 117 on butyrate.a

### 3.3 Proteomic responses per metabolic category

The 1772 proteins identified by MS-based proteomics were grouped into 27 metabolic categories using the Mercator program, which was conceived for plant sequences but can also be used for microalgae (May et al., 2008; Davidi et al., 2014), including *Polytomella* sp (Fuentes-Ramírez et al., 2021). Several other databases were used to confirm identities in case of doubt especially for the entries listed in Table I, as described in the Material and Methods section. Any redundant entries in the Mercator results were removed and retained in only one metabolic category. Of the 1772 statistically relevant proteins, 217 could not be annotated (12.2%) and 411 were annotated but could not be assigned to any Mercator category (23.2%) (Fig 3A). These percentages are similar or significantly lower than found in whole cell proteomic studies of other microalgae such as *C. reinhardtii* or *Dunaliella bardawil* (May et al., 2008; Davidi et al., 2014). A number of proteins that were not readily categorized but to which an EC number could be assigned, were included in the category ‘Enzyme classification’. The “protein biosynthesis” category is the metabolic category most represented with 209 proteins (11.8% of total proteins), followed by “protein homeostasis” (99 proteins; 5.6%) and “lipid metabolism” (84 proteins; 4.7%) (Fig. 3A). In addition, we calculated the cumulated iBAQ values to yield the total protein abundance per Mapman category in both conditions (Fig. 3B). The iBAQ value of a protein reflects its intra-sample abundance, and this calculation thus tries to estimate the impact of a condition on the accumulation of proteins related to a certain pathway. Among identified proteins, those in the categories “protein biosynthesis” and “cellular respiration” contribute most to the cumulative iBAQ value. Two major differences are seen between the two conditions. For acetate there is a more pronounced accumulation of proteins involved in protein biosynthesis (29% compared to 19.2% for butyrate), whereas for butyrate an enrichment is observed for proteins involved in lipid metabolism (11.8% vs 4.5% for acetate). It is noted that the two unassigned categories (annotated or not annotated) each contribute proportionally much less to the total iBAQ than to the number proteins, reflecting the fact that high abundance proteins are more likely to be attributed to a specific function.

**Table I.**
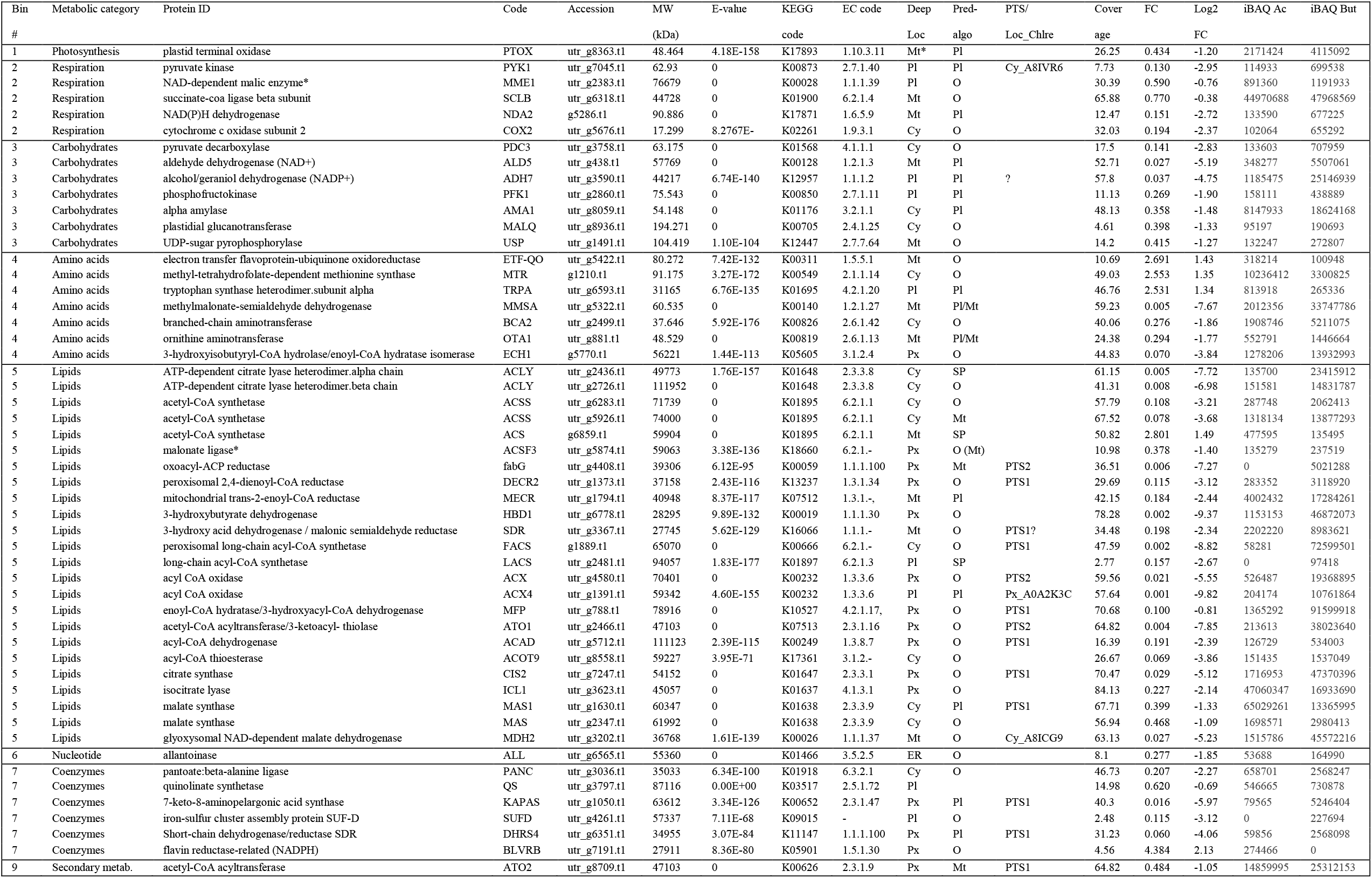
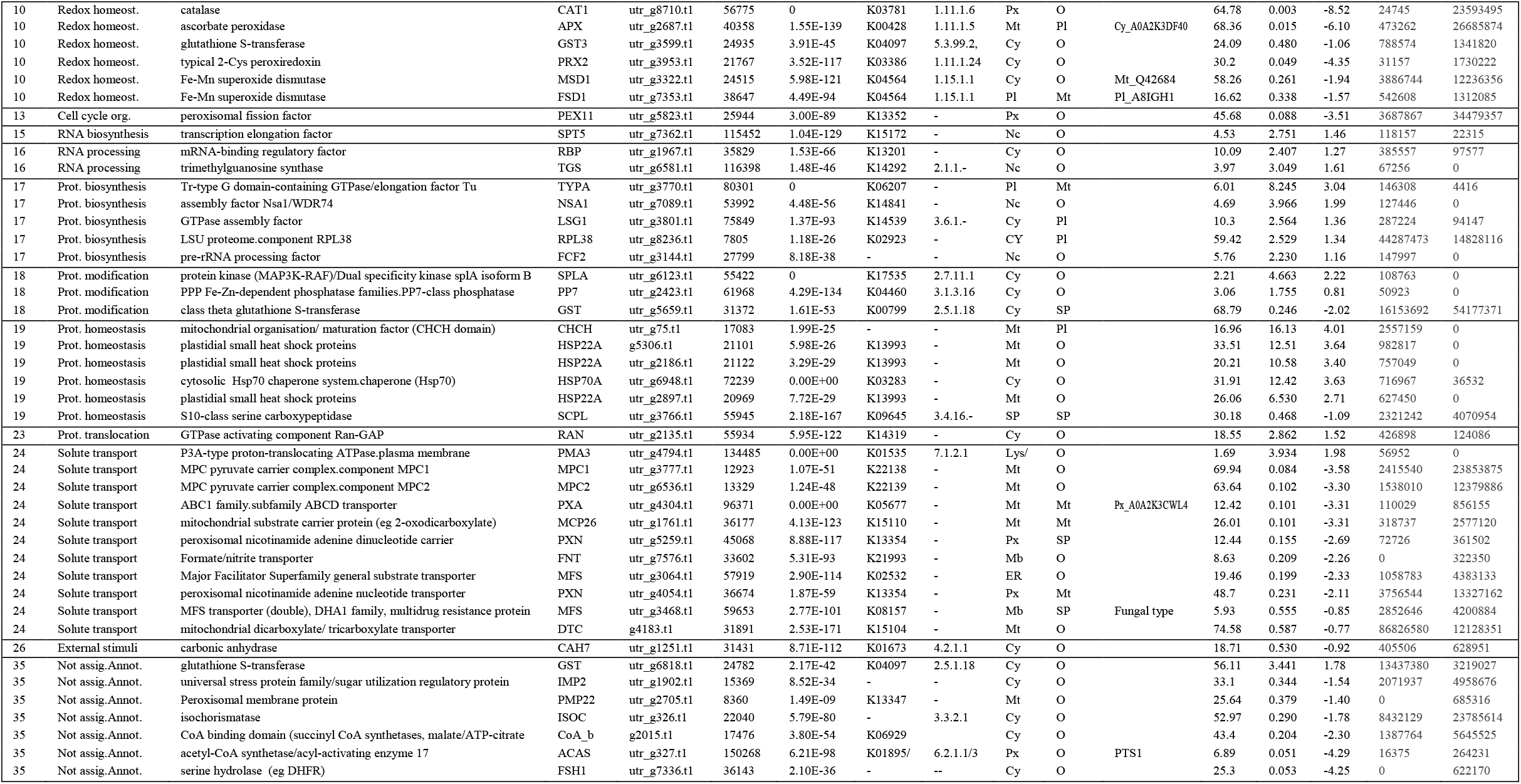
Overview of the most pertinent proteins with significant FoldChange values arranged by metabolic category as determined by Mercator. PTS, presence of peroxisome targeting signal; Loc_Chlre, location (when discordant) and accession no. of *C. reinhardtii* ortholog. Proteins corresponding to the E-values are given in the suppl. Table I.

**Figure 3.**
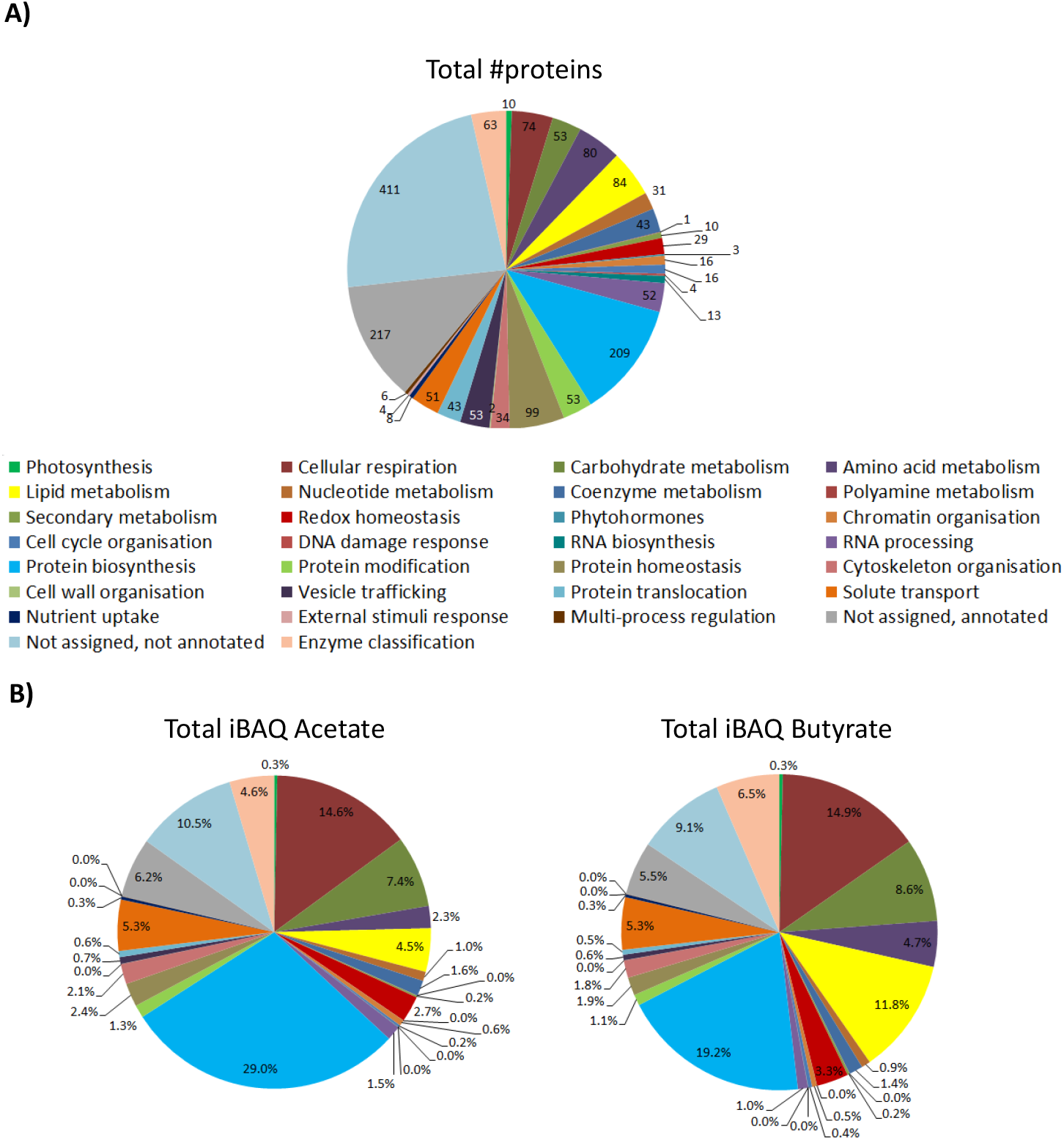
Overview of the *Polytomella* proteome revealed by the differential approach, represented per metabolic category as determined by Mercator. A) Total number of proteins identified in both acetate and butyrate-growing cells. B) Cumulative iBAQ values of total acetate and butyrate proteomes give an indication of the total protein abundance per category.

Focussing on proteins differentially-expressed (DE) between the acetate and butyrate conditions, the above global trend is confirmed but a more detailed picture emerges (Fig. 4). Relative to acetate, more proteins are overaccumulated on butyrate in the categories related to primary metabolism *i*.*e*. cellular respiration (4), carbohydrate metabolism (17), amino acid metabolism (10) and lipid metabolism (25), as well as the categories redox homeostasis (9), solute transport (10) and coenzyme metabolism (5). In contrast, acetate grown cells overaccumulate more proteins involved in homeostasis (6), biosynthesis (13) and protein translocation (2) as well as in RNA processing (4), processes that can be considered as necessary for general cellular maintenance. Comparing the proportion of FC proteins in each category to the proportion in these categories of the 1772 identified proteins using the Exact Fisher’s test, a significant enrichment in butyrate was obtained for the categories carbohydrate- and lipid metabolism, redox homeostasis and solute transport. For the acetate condition, the protein biosynthesis and protein homeostasis were found enriched. Overall, it is clear that the butyrate utilization mobilizes a higher number of proteins, suggesting a more profound metabolic response compared to acetate.

**Figure 4.**
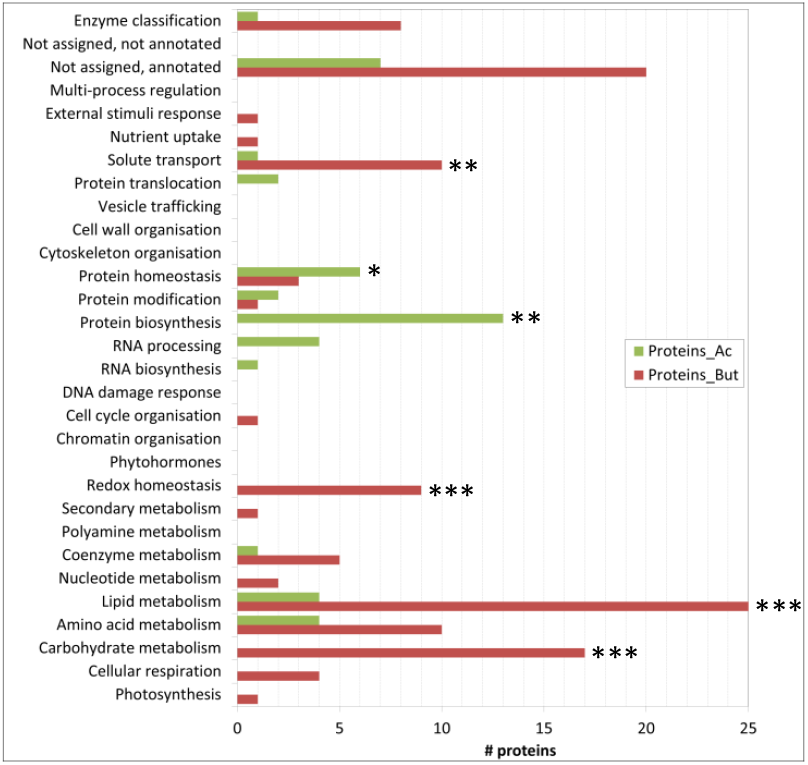
Focus on differentially expressed proteins, *i*.*e*. with a statistically significant difference in FC value. The number of proteins per category is given that are significantly more induced on either acetate or butyrate. Categories with a significant difference in FC proteins between either the acetate or the butyrate condition with respect to the background in the Fisher test are indicated with one asterisk (p-value<0.05), two asterisks (p<0.01) or three asterisks (p<0.001). Note that some categories do not contain any differentially expressed proteins, and the fact that some categories do not exhibit any FC proteins in either the acetate or butyrate condition does not mean there is no protein.

### 3.4 Metabolic pathways involved in butyrate response

To obtain a global metabolic representation of *Polytomella* sp., MapMan was used to project the FC values of individual proteins onto the different metabolic pathways. In section 3.5 these pathways will be discussed in detail. In the category lipid metabolism, the highest FC values are for enzymes of fatty acid synthesis, fatty acid degradation and the glyoxylate cycle (Fig. 5). The enzymes of the latter two functional groups are predicted to localize mostly to the peroxisomes, based on peroxisomal targeting prediction via software algorithms, manual analysis of the presence of peroxisomal targeting sequences PST1 and PST2, and/or the relatedness to peroxisomal enzymes in the green alga *C. reinhardtii* (Table I; see also suppl. Table I). Peroxisomes, organelles primarily dedicated to peroxide detoxification, seem to play a major role in butyrate assimilation in *Polytomella*. Butyrate being a fatty acid, the MapMan category “fatty acid degradation” is expected, and corresponds to what is described for other organisms (De Preter et al., 2012; De Meur et al., 2020). The glyoxylate cycle upregulation is in line with the fact that butyrate degradation leads to the production of 2 molecules of acetyl-CoA, which together with glyoxylate is the central entry point into the cycle. Fatty acid synthesis is also upregulated and is related in part to the activation with Coenzyme A, and seems to indicate cellular shift in the production sites and levels of acetyl-CoA (Pietrocola et al., 2015). Fatty-acid synthesis intermediates may for example be needed for the production of cofactors such as biotin and lipoic acid (Alban et al., 2000). Indeed, several enzymes involved in cofactors synthesis were found to be strongly upregulated as well. In addition, several enzymes (such as catalase) involved in antioxidant defense were strongly upregulated (Fig. 5), which can be directly or indirectly a consequence of the degradation of butyrate, notably by the formation of H_2_O_2_ at the site of the acyl-CoA oxidase (ACX) enzyme (Kato et al., 2021) but also due to the fact that butyrate is 25% more reduced compared to acetate, leading to an increased generation of reducing equivalents such as NADH.

**Figure 5.**
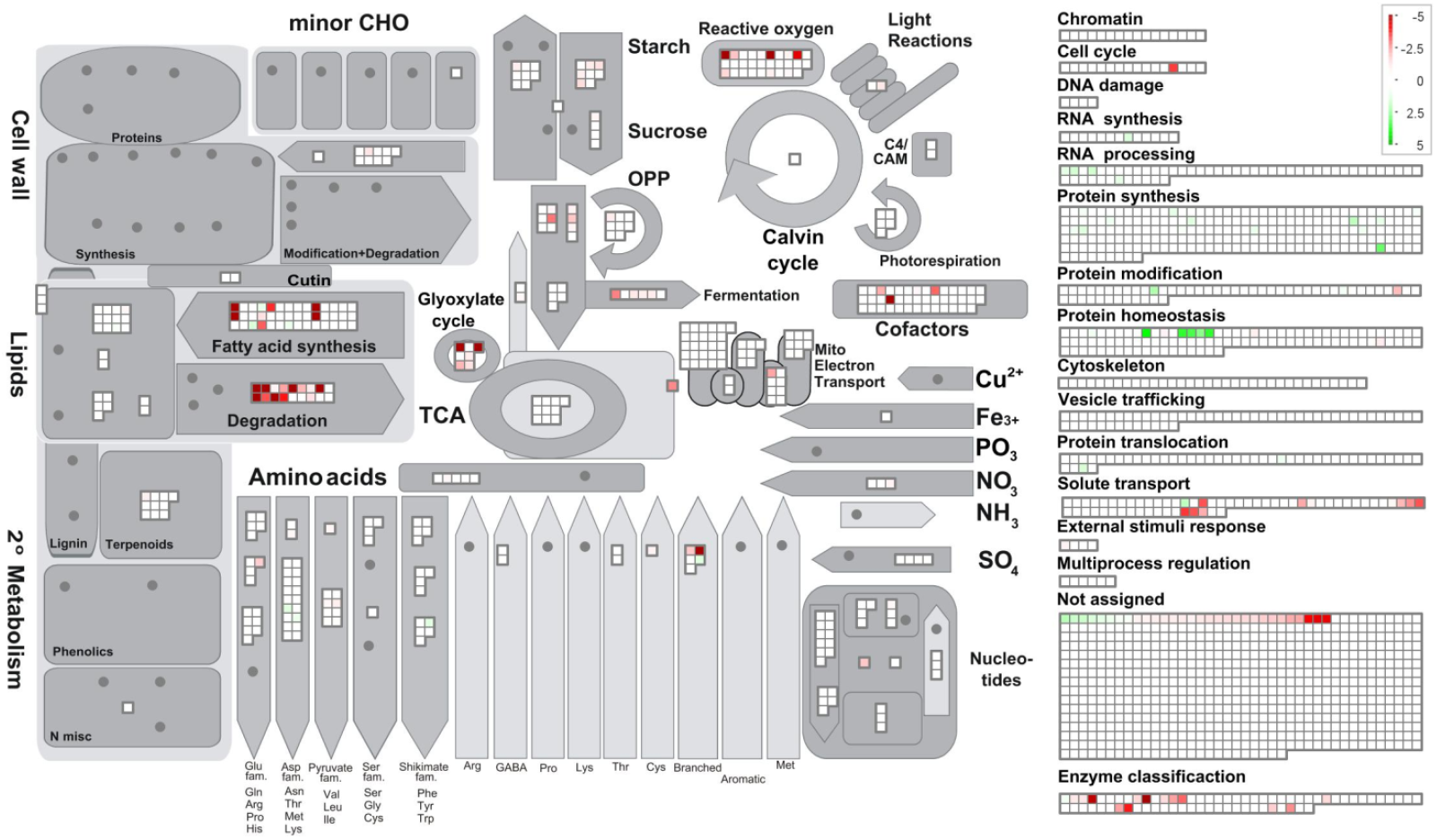
Schematic representation of cellular metabolism as a function of metabolic (sub)categories and log2 FoldChange values using the program MapMan. Metabolic categories were analyzed using the web program Mercator. For all entries with a FC score applies p-value <0.004. Note that for increased visibility of the lower range of the log2 FC scale was set from 5 to -5 whereas a few proteins on butyrate actually show higher FC values.

The amino acid metabolic pathway most affected by the C-source is the formation and subsequent degradation of branched chain amino acids (BCAAs). It appears to take place largely in the peroxisome based on the subcellular targeting predictions of the corresponding proteins, unlike in its close relative *C. reinhardtii* where it occurs in the mitochondria (Liang et al., 2019). It is noted that some of these enzymes are partitioned by MapMan into the lipid metabolism category. Carbohydrate metabolism, notably starch degradation, is upregulated on butyrate, reflected in the lower starch content in butyrate grown cells (Fig. 1D), which suggests an increased mobilization of glucose into pyruvate for downstream metabolic pathways. Finally, several peroxisomal and mitochondrial-type solute transporters are clearly induced on butyrate, which indicates increased exchange of a variety of metabolites between these compartments.

### 3.5 Reconstruction of the butyrate metabolic network

A list of those proteins that exhibit the most pronounced fold change sorted according to the Mercator categories is presented in Table I, alongside the most relevant data pertaining to protein databases and proteomic parameters. Protein identities were obtained by homology searches using Mercator and other programs for further validation (see material and methods), and their functional characteristics and their known or predicted intracellular localization were used to reconstruct the most important metabolic pathways in *Polytomella* sp. cells growing on butyrate relative to acetate. The most pronounced enrichment of enzymes involved in butyrate assimilation is found in the peroxisomes, small spherical organelles (0.2-1.5µm) that lack DNA and are surrounded by a single membrane that derives from the endoplasmic reticulum (Hu et al., 2012). Originally described as organelles that harbor oxidases that produce H_2_O_2_ and catalase for its detoxification (De Duve and Baudhuin, 1966), they are known to compartmentalize a large diversity of functions among which fatty acid β-oxidation (Gabaldón, 2010). In *Polytomella caeca*, small organelles distinct from mitochondria were identified that contained catalase activity (Gerhardt, 1971). Although previously typical peroxisomes could not be identified in *C. reinhardtii* (Silverberg, 1975), recently the β-oxidation enzyme acyl-CoA oxidase as well as catalase were identified in peroxisomes (Kong et al., 2017; Kato et al., 2021). Also, the enzymes of the glyoxylate cycle were shown in *C. reinhardtii* to localize to the peroxisome (or glyoxysome) (Lauersen et al., 2016). The proposed butyrate metabolic network is depicted in Fig. 6 and 7. Its operation depends not only on the presence of the appropriate enzymes, but also on their intracellular localization. Unfortunately, intracellular locales remain uncertain for some enzymes, due to lack of experimental evidence, possible N-terminal truncation of the gene models and ambiguous or obviously erroneous results from the prediction algorithms, which have not been trained specifically for *Polytomella*. Still, for most of the highly butyrate-induced proteins, reasonable deductions could be made that permit drawing a coherent picture of the intracellular location of the involved metabolic pathways (Table I).

**Figure 6.**
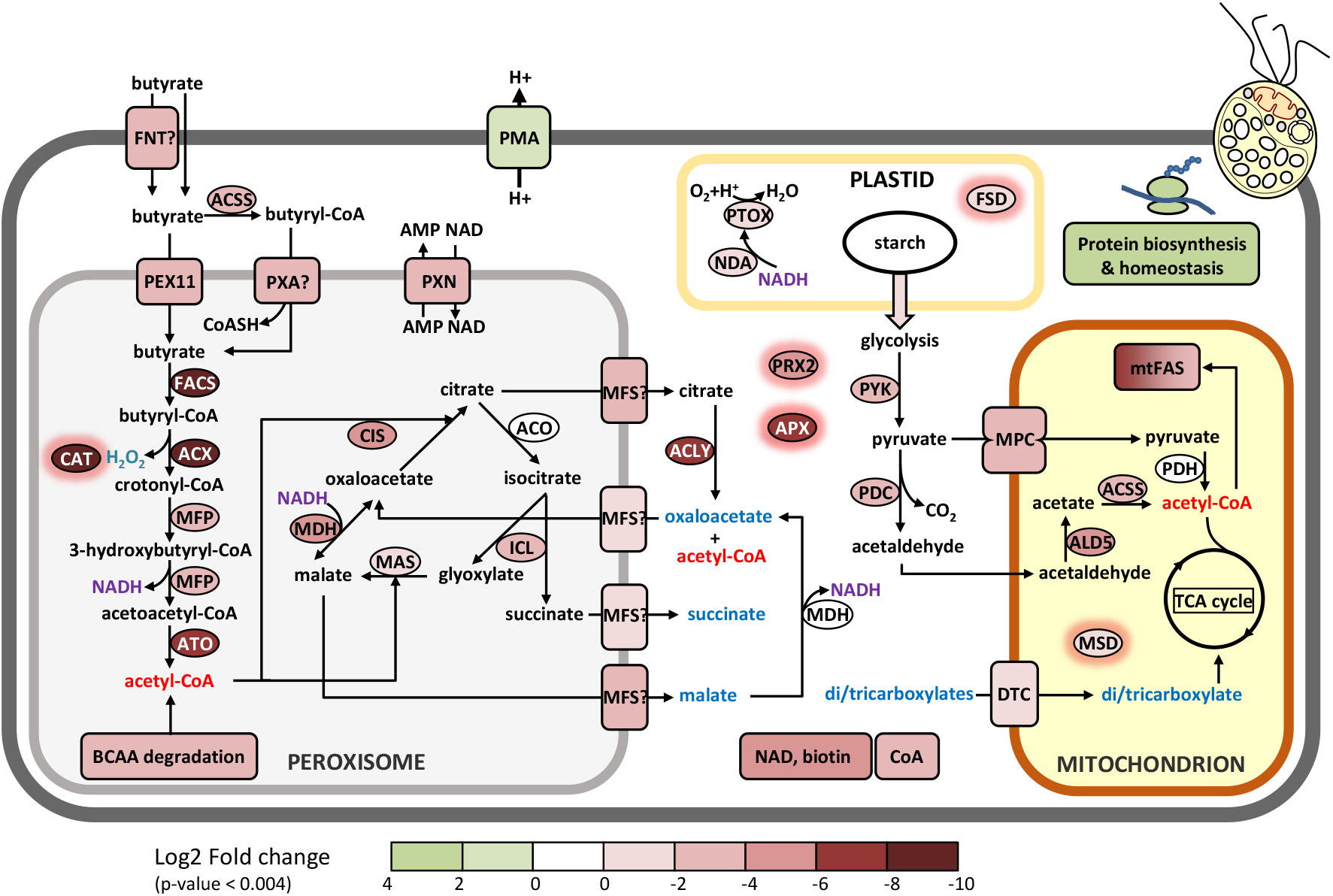
Proposed metabolic reconstruction of the assimilation pathway of butyrate in the peroxisome and interactions with other cell compartments. Taken into account are the FoldChange values and cellular localization based on software prediction (DeepLoc), manual verification of targeting signals and previous studies. The log2 FC acetate/butyrate value is indicated by the color codes. All di/tricarboxylic acids that may be imported into the mitochondria are indicated in blue. Enzymes with a red halo are involved in antioxidant defense. Enzymes codes and further information can be found in Table I and the Suppl. Table.

**Figure 7.**
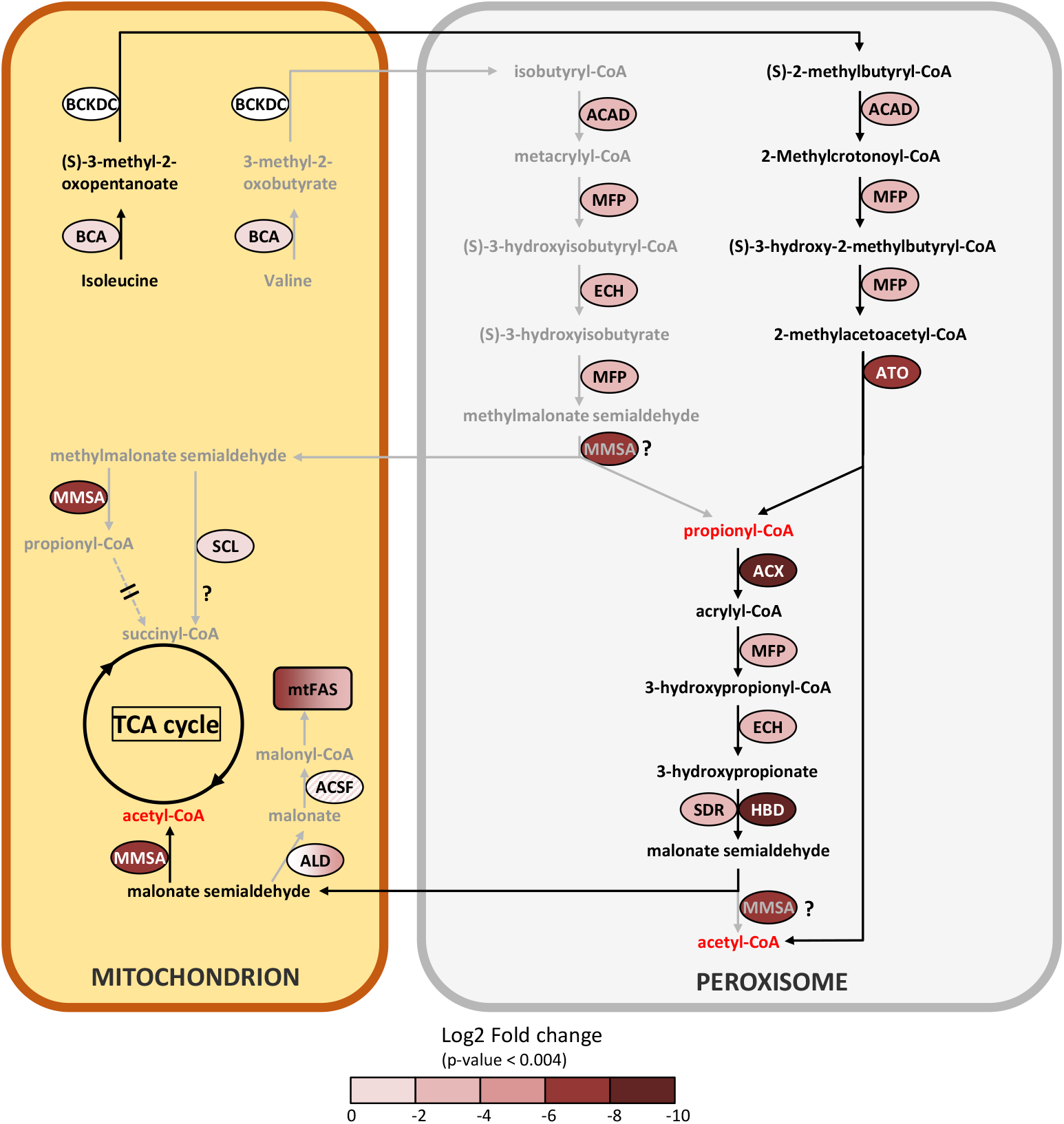
Metabolic reconstruction of the branched amino acid degradation pathway in the peroxisome and proposed interactions with the mitochondria. The log2 fold change But/Ac is indicated by the color codes. Further information can be found in Table I. Arrows in grey represent less likely or hypothetical pathways, proposed pathways use black arrows. ACSF exhibits an FC value with p>0.004. ALD color gradient indicates different isoforms with FC values between 0-6.

#### 3.5.1 Peroxisomal butyrate assimilation pathway

Once butyrate is taken up by the cell it must be activated to butyryl-CoA before it can enter the β-oxidation pathway in the peroxisome (Fig. 6). Candidates for this function were sought among the proteins most highly induced by butyrate. The highest induction level (FC 0.002; see Table I for Log2FC values) was found for g1889, which harbors at its C-terminus a typical PTS1 signal (Neuberger et al, 2003). It was annotated by Mercator as long-chain acyl-CoA synthase/ligase (LACS), an enzyme that does not act on fatty acids of less than 12 C-atoms (Wu et al., 2020). However, different types of fatty acyl CoA synthases can be found to be more closely related to the *Polytomella* enzyme using BLAST (45-50% ID), including medium chain acyl-CoA synthase and even bacterial 3-methylmercaptopropionyl-CoA ligases. We thus changed the annotation of the enzyme to fatty acyl-CoA synthase (FACS). This enzyme is the prime candidate for the production of butyryl-CoA in *Polytomella* sp., pending further experimental confirmation. A true LACS isoenzyme was also detected (UTR_g2481, FC 0.16), but since its expression level is much lower than FACS, this enzyme is unlikely to act as the first step of butyrate assimilation. Two acyl CoA oxidases (ACX) were also highly butyrate-specific, with a FC value of 0.001 for ACX4 and 0.021 for its paralog ACX. ACX4 was predicted to be localized in the plastid but homologs in other algae including *C. reinhardtii* retrieved by BLAST searches (∼51% ID) are annotated as peroxisomal (Table I). Examination of the N-terminus of isoform ACX reveals a typical PTS2 signal and shows 34% sequence identity to *C. reinhardtii* ACX2, which catalyzes the first step of peroxisomal fatty acid β-oxidation in the green alga *C. reinhardtii* (Kong et al. 2017). These two ACX enzymes likely oxidize butyryl-CoA into crotonyl-CoA, which is converted into 3-hydroxybutyryl-CoA and then into acetoacetyl-CoA by respectively the enoyl-CoA hydratase and the hydroxyacyl-CoA dehydrogenase activities of the multifunctional protein MFP (Fig. 6). Compared to ACX4, MFP was found to be more modestly induced by butyrate (FC 0.1). The final step of β-oxidation is carried out by acetoacetyl-CoA thiolase (ATO1, FC 0.004), converting acetoacetyl-CoA into two molecules of acetyl-CoA.

All 4 identified enzymes have typical PTS peroxisome targeting signals (Table I) and are thus confirmed to be peroxisomally targeted. The fatty acid β-oxidation pathway used by *Polytomella* sp. is typical for algae and plants, where it is found in both mitochondria and peroxisomes (Kong et al., 2017; Li-Beisson et al., 2019; Pan et al., 2020; Kato et al., 2021).

We currently have no data on a mitochondrial β-oxidation pathway in *Polytomella* sp., but our study clearly indicates that the peroxisomal pathway is key in the assimilation of butyrate. The peroxisomal β-oxidation pathway of butyrate in *Polytomella* sp. differs from that in non-photosynthetic organisms and bacteria by the CoA activation step and by the presence of MFP. Besides the use of a FACS type enzyme, CoA activation of butyrate in mammals and microorganisms and can also use a butyrate-CoA ligase/synthetase (EC 6.2.1.2) (De Preter et al., 2012). Conversely, fermentative butyrate production from butyryl-CoA in bacteria mainly occurs via phosphate butyryltransferase (EC 2.3.1.19) + butyrate kinase (EC 2.7.2.7) (Walter et al., 1993) or butyryl-CoA:acetate CoA-transferase (EC 2.8.3.8) (Duncan et al., 2002). The presence of a bifunctional MFP is the hallmark of peroxisomal β-oxidation in plants, fungi and microalgae (Arent et al., 2010). In contrast, its two reactions are carried out by separate enzymes in mitochondrial β-oxidation of butyrate in mammalian colonocytes (De Preter et al., 2012).

#### 3.5.2 Glyoxylate cycle and citrate/malate shuttles

The glyoxylate cycle comprises five enzymes which are mostly present in peroxisomes but some, depending on the organism, can also be found in the cytosol, possibly to protect them from ROS-induced damage. In *C. reinhardtii*, the glyoxylate cycle allows growth on acetate (Lauersen et al., 2016) following its conversion into acetyl-CoA. Glyoxylate and acetyl-CoA are converted into malate and further into citrate and succinate, which are exported to enter central carbon metabolism, and can replenish the pool of TCA cycle intermediates. In *Polytomella*, the typical glyoxylate cycle enzymes malate synthase (MAS1, FC 0.4) and isocitrate lyase (ICL1, FC 0.23) are induced on butyrate, but at a lower level than citrate synthase (CIS2, FC 0.03) and malate dehydrogenase (MDH2, FC 0.03) (Fig. 6). Aconitase (ACO) is also part of the cycle and three isoforms were detected (suppl. Table I) but none were induced significantly by butyrate. MAS1 is confirmed to localize in the peroxisome based on the presence of a typical PTS1 targeting signal at its C-terminus (Gonzalez et al., 2011). CIS2 is predicted to be targeted to the peroxisome by DeepLoc (Almagro Armenteros et al., 2017) (Table I) and its closest homologs are found in peroxisomes in plants and yeast (Kunze et al., 2006; Rottensteiner and Theodoulou, 2006). *Polytomella* CIS2 and MDH2 show highest amino acid sequence identities to peroxisomal/glyoxysomal-type homologs. ACO may also be peroxisomal since three out of four amino acids of the consensus PTS2 signal are present (RV-X5-RA instead of RV-X5-H/QA) (Gonzalez et al., 2011). In *C. reinhardtii*, only ICL was found in the cytosol with the other 4 enzymes in peroxisomal microbodies (Lauersen et al., 2016). Earlier it was found that in *Polytomella caeca*, microbodies separated from mitochondria on a sucrose gradient contained MAS and a minor part of ICL activity, with most of it being cytosolic (Haigh and Beevers, 1964). Since these authors concluded that the ICL activity within the microbodies accounted for the observed acetate assimilation, it may be assumed that ICL functions in the peroxisome, as is also predicted by DeepLoc (Table I). The upregulation of ICL1 on butyrate with respect to acetate may indicate an increased activity of the cycle, resulting in higher glyoxylate and succinate production. While glyoxylate serves to produce malate via MAS in the peroxisome, succinate may be exported to the cytosol and further imported into the mitochondria for use in the TCA cycle (Fig 6). The pronounced increase of MDH2 may be related to the production of NADH by MFP during butyrate β-oxidation in the peroxisome. At the expense of NADH, MDH2 can convert oxaloacetate (OAA) into malate, which can be later exported to the cytosol (Fig. 6) via a malate/OAA transporter (Rottensteiner and Theodoulou, 2006). Export of citrate produced by CIS2 from oxaloacetate is likely occurring under butyrate growth considering the very strong induction of the cytosolic ATP:citrate lyase (ACLY, FC 0.005). This enzyme produces acetyl-CoA and oxaloacetate from citrate in the cytosol, thus allowing the export of acetyl-CoA. The oxaloacetate resulting from ACLY can be re-imported to replenish the peroxisomal oxaloacetate pool for the proper functioning of the glyoxylate cycle (Fig. 6). Despite the significant relatedness of MDH2 to its peroxisomal homolog in *C. reinhardtii*, (62% amino acid identity) it can currently not be excluded that MDH2 is cytosolic, and if so, this would indicate an increased need for malate/oxaloacetate shuttle activity to sustain increased production and export of citrate from the peroxisome. A cytosolic localization for MDH, ACO and ICL occurs in yeast and does not impact glyoxylate cycle function (Rottensteiner and Theodoulou, 2006). The fact that MDH2 and CIS2 are more induced than ICL1 and MAS1 points to an apparent increased need for a malate/oxaloacetate shuttle and citrate export in butyrate metabolism. With regard to CIS, it was shown that during ^14^C-acetate assimilation of *P. caeca*, malate was by far the most important immediate product incorporating ^14^C, with succinate and (mitochondrial) fumarate 10-fold lower, but hardly any citrate was formed (Haigh and Beevers, 1964). Conversely, judged from the upregulation of CIS2 and ACLY, butyrate seems to specifically induce peroxisomal citrate production. Since the outcome of butyrate utilization is the increased production of acetyl-CoA in the cytosol from ACLY, an induction of fatty acid synthesis may be expected in organelles. It is interesting that in human colonocytes, butyrate stimulates cell proliferation via histone acetylation in the nucleus, involving the production of acetyl-CoA by ACLY (Donohoe et al., 2012). In our study, no histone acetyltransferase was found differentially expressed.

#### 3.5.3 Transporters and metabolite exchange

Relative to acetate, butyrate induced 10 membrane bound transporter proteins with significant associated FC values (Table I, category “solute transport”), among which genuine peroxisomal transporters. However, the identity of the proteins transporting butyrate or butyryl-CoA into the peroxisomes remains uncertain. The ABCD transporter PXA (FC 0.1) shows high similarity to an ABCD transporter in *C. reinhardtii* (A0A2K3CWL4/Cre15.g637761) which was confirmed to be involved in the import of activated long-chain fatty acids from the cytosol to the peroxisomal matrix, similar to the yeast peroxisomal ABC transporters PXA1 and PXA2 (Hettema et al., 1996). PXA targets long-chain fatty acyl-CoA molecules, which butyryl-CoA is not, so its involvement in the import of butyryl-CoA from the cytosol is uncertain. It might be necessary instead to channel CoA into the peroxisome for CoA homeostasis. Potentially, the two cytosolic acetyl-CoA synthase-type enzymes (ACSS) that were upregulated by butyrate (FC 0.08, 0.11) may provide substrates for this process. Another induced transporter belongs to the PEX11 family (FC 0.09, category cell cycle in Table I), a membrane protein that promotes peroxisome division in eukaryotes and is crucial for medium-chain fatty acid (MCFA) beta-oxidation. It was proposed that in yeast, PEX11 provides MCFAs including butyrate to the peroxisome interior for CoA activation, effectively fulfilling a transporter function (van Roermund et al., 2000). Two further proteins show clear homology to the peroxisomal NAD carrier PXN (FC 0.155, 0.231) in *Arabidopsis thaliana*, which mediate the import of NAD into peroxisomes against AMP (van Roermund et al., 2016). PXN belongs to the mitochondrial carrier (TC 2.A.29) family, which contains also peroxisomal transporters. Butyrate may thus increase the import of cofactors in the peroxisome, possibly linked to enhanced peroxisome biogenesis.

Three proteins were identified as mitochondrial transporters, indicating also an involvement of mitochondrial metabolism in butyrate utilization. Two of them are subunits of the mitochondrial pyruvate transporter (MPC1, FC 0.08 & MPC2, FC 0.1), an oligomeric complex of approximately 150 kDa in the inner mitochondrial membrane which constitute the sole entry point into the mitochondria of pyruvate, produced by glycolysis or from malate. The upregulation of this carrier is of fundamental importance in establishing the metabolic programming of a cell (Bricker et al., 2012). Once in the matrix, pyruvate can be converted into acetyl-CoA by the pyruvate dehydrogenase complex (PDH) and feed the TCA cycle (McCommis and Finck, 2015). Cycle turnover produces CO_2_ and reducing power further used for ATP production, but intermediates can be siphoned off such as citrate, which can exit the mitochondria and be cleaved back to acetyl-CoA and oxaloacetate by ATP citrate lyase (ACLY) in the cytosol. Another transporter identified is homologous to the mitochondrial dicarboxylate/tricarboxylate transporter DTC (FC 0.6) in *A. thaliana* (Millar and Heazlewood, 2003). DTCs transport dicarboxylic acids (eg malate, oxaloacetate) and tricarboxylic acids (eg citrate, isocitrate) into the mitochondrial matrix. In view of the FC value of 0.6, the role of DTC is only modestly increased in butyrate metabolism, but the iBAQ value for DTC is highest among transporters and represents one of the most abundant among butyrate induced proteins, illustrating the importance of di/tricarboxylates for mitochondrial metabolism. Seeing that the levels of the proteins of mitochondrial respiration (OXPHOS complexes) are not induced by butyrate, it can be proposed that increased import of pyruvate and dicarboxylates feeds other metabolic pathways, such as amino acid synthesis (see below).

Furthermore, two general substrate transporters of the Major Facilitator Superfamily were identified (MFS, FC 0.2, 0.56). MFS transporters can only transport small solutes in response to chemiosmotic ion gradients and can be found anywhere in the cell, so their proposed placement in the peroxisomal membrane for transport of carboxylates is speculative (Fig 6). The monocarboxylate transporters MCT that are known in humans to transport acetate and butyrate across the plasma membrane are also members of the MFS family (Casal and Leão, 1995), but they are not orthologous to the two MFS proteins mentioned above. Also, a typical MCT could not be found in the *Polytomella* sp. genome. However, a member of the formate/nitrite transporter (TC 2.A.44) family (FNT, FC 0.21) was found to be induced by butyrate. FNT transporters transport monovalent anions and are not strictly selective as they can use nitrite or formate but also larger organic anions such as lactate and acetate (Lu et al., 2012). It may thus be that this FNT is actually responsible for butyrate transport over the plasma membrane in *Polytomella* sp. Further biochemical and genetic studies need to confirm whether this is the case. Five genes belonging to the GPR1/FUN34/YaaH (GFY) superfamily and homologous to bacterial acetate/succinate channels were found to be induced by acetate in *C. reinhardtii* and possibly implicated in intracellular acetate transport (Durante et al., 2019). In the *Polytomella* sp. genome, 4 GFY genes were identified of which only 1 (UTR_1663.t1) was found in the proteome induced by butyrate at FC 0.636 (but with p>0.004). One transport protein is actually slightly downregulated, a P-type plasma membrane H+-ATPase (PMA3, FC 3.9) that exports cellular protons (Morth et al., 2011). In *C. reinhardtii*, increased PMA expression was found to improve tolerance to high CO_2_ concentrations, which are toxic due to the acidification of the cell interior (Choi et al., 2021). Acetate or butyrate are imported in the protonated form and dissociate in the cytosol. It can be hypothesized that butyrate necessitates less expulsion of H^+^ since it contains relatively fewer carboxylic acid groups at only 1 COOH per 4 C-atoms while acetate contains 1 COOH per 2 C-atoms.

#### 3.5.4 Antioxidant defense

Reactive oxygen species (ROS) are important in cellular signaling, but stress conditions may cause increased ROS production and result in oxidative damage (Rezayian et al., 2019). Various antioxidant defense systems to neutralize ROS exist in different cellular compartments, especially in organelles that are major sources of ROS such as H_2_O_2_ (Roy et al., 2021). The butyrate metabolic responses include a total of 10 proteins that are associated to antioxidant defense. One catalase isoform, co-orthologous to the *C. reinhardtii* CAT1 isoform that localizes to peroxisomes (Kato et al., 2021) and similarly endowed with a non-canonical C-terminal PTS1 signal (S*V*L), was found markedly induced on butyrate (FC 0.003), while a second CAT1 ortholog was far less induced. No clear ortholog was found for the *C. reinhardtii* ER-localized CAT2. Catalase upregulation probably relates to the β-oxidation of butyrate, where it allows the detoxification of H_2_O_2_ produced by ACX into H_2_O and O_2_. Using density gradients, (Gerhardt, 1971) found the catalase and malate synthase activities in different particulate fractions in *P. caeca*, which raised the question whether different types of peroxisomes exist in the cell. A study to detect peroxisomes using CAT-specific staining seemed to shown that indeed different staining intensities existed in a sample of isolated peroxisomes (Gerhardt and Berger, 1971).

Ascorbate peroxidase (APX, FC 0.015) was highly induced by butyrate. It is part of the glutathione-ascorbate cycle (GAC) that uses electrons from NAD(P)H for the reduction of H_2_O_2_ to H_2_O. Other GAC enzymes were also induced on butyrate, such as glutathione S-transferase (GST3, FC 0.51), while glutathione reductase (GR) was not. The GAC can be found in different cellular compartments such as plastids, mitochondria, peroxisomes and the cytosol, (Caverzan et al., 2012). Since no coherent targeting signals were found for *Polytomella* APX, GST3 and GR, the GAC was tentatively placed in the cytosol in Fig. 6 since the closest homologs of these enzymes in *C. reinhardtii* are predicted to be cytosolic. A typical 2-Cys peroxiredoxin (PRX2, FC 0.05) was found highly induced on butyrate. PRXs are important for cellular redox signaling and antioxidant defense as they detoxify organic hydroperoxides (R-O-OH) that can be formed from the reaction of organic molecules with H_2_O_2_ (Liebthal et al., 2018). Another important ROS scavenging enzyme is superoxide dismutase, which produces H_2_O_2_ from O_2_^-^. Two SOD isoforms were found, only modestly induced on butyrate (MSD1, FSD1, FC 0.26, 0.34).The localization of PRX2 and M/FSD1 is uncertain as they lack typical targeting signal and were predicted to be cytosolic by DeepLoc. Intriguingly, the levels of plastid type alternative oxidase (PTOX, FC 0.43), which functions in plastid redox homeostasis by oxidizing plastoquinol to reduce O_2_ to H_2_O in photosynthetic organisms (Krieger-Liszkay and Feilke, 2016), and the NAD(P)H dehydrogenase that feeds electrons into the plastoquinone pool (NDA2, FC 0.15), were higher on butyrate than acetate. This suggests a role of chlororespiration in antioxidant defense in *Polytomella*. It is of note that these enzymes are found in the thylakoid membrane in *C. reinhardtii*, while in *Polytomella* sp. a localization to the amyloplast envelope is proposed in absence of a description of any intraplastidial membrane system (Fuentes-Ramírez et al., 2021).

Interestingly, photosynthesis in rice leaves was found to be protected due to a more efficient antioxidant response when CAT and APX activity were limited, which was proposed to be because of higher H_2_O_2_ levels that exert a positive regulatory influence (Sousa et al., 2019). First off, it would be interesting to know whether the C4 fatty acid butyrate does in fact induce β-oxidation in photosynthetic algae since it is known that *C. reinhardtii* favors membrane turnover over β-oxidation in presence of exogenous C16 fatty acid (Kato et al., 2021). If not it would explain directly why butyrate is poorly used by *C. reinhardtii* (Lacroux et al., 2020). If butyrate does enter β-oxidation, increased CAT and APX could modify ROS levels and, in the view of (Sousa et al., 2019), interfere with photosynthesis and hinder the growth on butyrate by green algae, as observed by (Lacroux et al., 2020). *Polytomella* sp does not perform photosynthesis and is thus less affected by the very strong antioxidant responses elicited by that the β-oxidation of butyrate, which is possibly at the basis of its capacity to grow well on butyrate.

#### 3.5.5 Branched chain amino acid degradation

A number of proteins induced on butyrate, placed in both the lipid and amino acid metabolism categories (Table I), potentially participate in the degradation of branched chain amino acids (BCAAs), by which valine is converted to propionyl-CoA, while degradation of isoleucine produces both propionyl-CoA and acetyl-CoA (Fig. 7). This BCAA pathway is described in prokaryotes and in eukaryotes such as mammals, yeasts and plants (Binder, 2010), but also in microalgae, for example the diatom *Phaeodactylum tricornutum* (Pan et al., 2017) or *C. reinhardtii* (Liang et al., 2019). The first enzyme of this pathway is branched-chain aminotransferase (BCA2, FC 0.28), probably located in the mitochondria. Except for the second enzyme, branched-chain alpha-keto acid dehydrogenase (BCKDC, FC 2.1), all enzymes in this pathway are induced by butyrate, in particular the central enzyme methylmalonate semialdehyde dehydrogenase (MMSA, FC 0.005). Most of the dowsntream enzymes seem to be targeted to the peroxisome in *Polytomella* sp. (Table I), recapitulating the situation in plants where BCAA degradation starts in the mitochondrion to yield CoA-esterified metabolites that are further converted in the peroxisome (Linka and Theodoulou, 2013).

Questions remain about this pathway in *Polytomella* sp. The pathway depends likely on the cellular localization of MMSA. While the *C. reinhardtii* methylmalonate semialdehyde dehydrogenase (ALDH6) is predicted to be mitochondrial, the *Polytomella* ortholog is predicted to be peroxisomal (Fig. 7). If the latter is the case, the degradation of both isoleucine and valine towards acetyl-CoA can occur unimpeded in the peroxisome. If we however entertain the possibility that MMSA is mitochondrial in *Polytomella* sp., the situation is different. MMSA is able to convert MMS into propionyl-CoA but also malonate semialdehyde (MS) into acetyl-CoA, so the degradation of isoleucine into MS (via propionyl-CoA) does not demand the presence of MMSA in the peroxisome when MS is imported into mitochondria. Mitochondrial MMSA can convert MS into acetyl-CoA which can be further utilized without problems. In the case of valine degradation, the absence of MMSA in the peroxisome would block the pathway at the level of the conversion of MMSA to propionyl-CoA, and in that case MMS would have to be exported to the mitochondria. Here, MMSA would convert MMS into propionyl-CoA, but its fate in the mitochondria would not be obvious: the enzymes necessary for its conversion to succinyl-CoA (propionyl-CoA carboxylase producing (S)-methylmalonate-CoA, methylmalonyl-CoA epimerase producing (R)-methylmalonate-CoA and methylmalonyl-CoA mutase to convert it to succinyl-CoA) could not be identified in the *Polytomella* genome. This is similar to the case of *P. tricornutum* where the epimerase step was not detected (Pan et al., 2017). In *P. caeca* cells grown on propionate, propionyl-CoA carboxylase activity could not be detected (Lloyd et al., 1968). The same is true in plants, unlike in mammals and bacteria (Linka and Theodoulou, 2013). The TCA cycle enzyme succinyl-CoA ligase (SCL, FC 0.77 at p>0.003), somewhat induced by butyrate, was found to be a promiscuous enzyme that produces also thioesters of malate, fumarate and glutarate among others (Nolte et al., 2014). It may be hypothesized that SCL can produce succinyl-CoA directly from methylmalonate semialdehyde (MMS). Alternatively, MMS may also be imported into mitochondria and converted into malonate by an aldehyde dehydrogenase (ALD5 EC:1.2.1.-; at least 5 enzymes are found in the proteome, suppl. Table I) and then further into malonyl-CoA via malonate ligase (ACSF3, FC 0.38 but with p>0.004), which serves as precursor for fatty acid synthesis (see 3.5.6). It is noted that even if ALD5/ACSF3 are not located in the mitochondria, the products can be imported into the organelle. Alternatively, the BCAA degradation pathway may be streamlined when MMSA is dually targeted to both peroxisome and mitochondria, which is known for other enzymes such as CAT (Petrova et al., 2004).

In the non-sulfur purple bacterium *Rhodospirillum rubrum*, which does not possess a glyoxylate cycle, the BCAA degradation pathway was identified as assimilatory during growth on butyrate (De Meur et al., 2020). Interestingly, *R. rubrum* grew 3-fold faster in presence of HCO3^-^, which serves as electron sink and helps antioxidant defense. However, since *Polytomella* sp. does possess an active glyoxylate cycle, the purpose of the BCAA degradation pathway in butyrate assimilation is not directly obvious. It could serve to produce propionyl-CoA for metabolic pathways such as the synthesis of coenzyme A. Leucine degradation has been found in *A. thaliana* alongside starch and lipid degradation in response to stress conditions that perturb cellular energy balance, such as senescence and carbon deprivation (Mentzen et al., 2008). In these conditions, it is conceivable that the cell makes up for a lack of energy and carbon by mobilizing internal reservoirs of sugar, lipid and amino acids. In *Polytomella*, all three are observed in cells growing on butyrate (see 3.5.6), whereby sugars and lipids may serve to produce amino acids such as BCAAs. We propose that butyrate is metabolized and yields acetyl-CoA and further di/tricarboxylic acids at a slower rate than acetate, which is compensated for by the degradation of BCAAs to produce acetyl-CoA and possibly succinyl-CoA that feed into carbon metabolism. A factor in the induction of the BCAA degradation pathway specifically may be the fact that it shares several enzymes with the β-oxidation of butyrate (ACX, MFP, ATO).

#### 3.5.6 Catabolic production and role of pyruvate

Pyruvate is at the crossroads of many metabolic pathways and is important in all cells and cell compartments (Shtaida et al., 2015). There are multiple indications that butyrate metabolism goes also through pyruvate, while it is not predicted to be directly involved in acetate utilization. First, most enzymes of glycolysis are upregulated several fold under butyrate, including pyruvate kinase (PYK1, FC 0.13) (Table I), which should result in increased pyruvate production and ATP. Compared to acetate, the balance seems to be shifted towards starch and glucose degradation, in line with the cellular sugar content being lower on butyrate (Fig. 1D) and the induction of the mitochondrial MPC transporter for pyruvate (see 3.5.3). This may not be leading to higher TCA cycle and OXPHOS activities since the necessary proteins are not induced, but pyruvate may instead be converted into amino acids such as alanine, and fatty acids (see below). Also, pyruvate decarboxylase (PDC3, FC 0.14) was induced, which should lead to acetaldehyde production in the cytosol. This could diffuse through the mitochondrial membrane and then be converted into acetate by NAD+ aldehyde dehydrogenase (ALD5, FC 0.03) and then acetyl-CoA by acetyl-CoA synthase (ACSS, FC 0.01). This scenario is supported by the fact that *Polytomella caeca* can grow on acetaldehyde (1mM) as sole external carbon source (Wise, 1968). It cannot be excluded that acetaldehyde is converted into acetate in the cytosol which is then imported into the mitochondria (see 3.5.8). Finally, there may also be a contribution from NADP malic enzyme (ME, FC 0.82 with p>0.004) converting malate to pyruvate in the cytosol. Since this enzyme was predicted to be targeted to the plastid, it can provide a source of pyruvate from imported malate. Via an MDH type enzyme (5 different MDH were detected in the proteome, suppl. Table I) in the plastid that produces NADH, malate may also feed the PTOX enzyme that was found induced on butyrate (see 3.5.4) and is implicated in maintaining cellular redox balance. Indeed, 2 MDH enzymes and PTOX were detected in a proteomics study of isolated non photosynthetic plastids from *Polytomella parva* (Fuentes-Ramírez et al., 2021). Pyruvate is also at the basis for the production of coenzyme A and NAD+, with pantoate:beta-alanine ligase (PANC, FC 0.21) of the CoA/ACP synthesis pathway and quinolinate synthase (QS, FC 0.62) of the NAD+ synthesis increased, suggesting a need of CoA/ACP in butyrate metabolism.

A clear upregulation is found of two of the four enzymes of the type II fatty acid synthesis (FAS) system, which uses an acyl carrier protein (ACP): 3-oxoacyl-ACP reductase (fabG, FC 0.006) and enoyl-ACP reductase (MECR, FC 0.184). In *C. reinhardtii*, the four different subunits of the type II FAS system are predicted to be dually targeted to the mitochondrion and chloroplast, similar to plants (Riekhof et al., 2005; Li-Beisson et al., 2013). Fatty acids produced in the plastid are used for the production of membranes, storage and signaling lipids (Li-Beisson et al., 2015). The FAs are unlikely to be destined for the production of storage lipids since levels are actually lower in butyrate grown cells (Fig. 1D). Fatty acids produced by mitochondrial (mt)FAS play various roles. The mtFAS pathway fuels the production of acyl-ACPs including octanoyl-ACP, which is a precursor of lipoic acid, a cofactor of several metabolic enzymes: pyruvate dehydrogenase (PDH), α -ketoglutarate dehydrogenase (KGDH), branched-chain α -ketoacid dehydrogenase (BCKDH), the glycine decarboxylase complex (GDC), and plastidial pyruvate dehydrogenase (ptPDH) (Guan et al., 2020). Four out of five of these enzymes are indeed upregulated on butyrate with FC values of 0.58-0.87, and although these values are not statistically sound enough to warrant inclusion in Table I, it does represents a clear trend. Paradoxically, the only enzyme not changed is BCKDH, which is involved in the BCAA degradation pathway that is highly expressed on butyrate. This suggests that the pathway is not regulated at the level of this enzyme, which is not an uncommon observation in biochemical studies (eg Nogaj et al., 2005). The mtFAS system is possibly induced to provide acyl-ACPs to two enzymes involved in biotin synthesis, 3-oxoacyl-ACP reductase (OAR, FC 0.06 and 7-keto-8-aminopelargonic acid synthase (KAPAS, FC 0.016), which uses pimeloyl-ACP as substrate.

Biotin is known to be a cofactor of certain carboxylase enzymes, including acetyl-CoA carboxylase (ACC) which functions 2 steps upstream of the FAS system producing malonyl-CoA from acetyl-CoA. ACC was the only enzyme with a biotin cofactor that was detected in the *Polytomella* proteome, and although it is not induced by butyrate it may be regulated post-translationally. It is noted that ACLY in the cytosol is strongly induced (see 3.5.2) and produces the acetyl-CoA that is a direct substrate for ACC. Biotin has been described to exert regulatory influences in cell signaling, for example the upregulation of glucose metabolism (Dakshinamurti, 2005), which was indeed induced on butyrate.

#### 3.5.7 Metabolic activities on acetate

The differential approach revealed that the proteins more abundant on acetate than on butyrate tend to relate to the protein biosynthesis and homeostasis (further referred to as proteostasis) rather than to specific metabolic pathways. Among the over-represented categories are “RNA processing” and “Protein biosynthesis, modification and homeostasis”. This includes heat-shock proteins, protein kinases, maturation-, elongation- and assembly factors. Although butyrate induced more proteins compared to acetate, there may also be downregulation signals produced in response to specific metabolic needs imposed by butyrate. A specific perception of an acetate-linked metabolite and associated signal cascades may also be involved. Some of the proteins suppressed by butyrate may suggest the implication of a mitogen-activated protein kinase (MAPKs) signal transduction pathway, which modulates important cellular processes such as proliferation, stress responses, apoptosis and immune defense via consecutive protein phosphorylations by serine and threonine protein kinases (Soares-Silva et al., 2016).

We note the induction of a protein of uncertain function, being either protein kinase (MAP3K-RAF) or dual specificity kinase splA isoform B (FC 4.663). It may suggest activation of a signal transduction pathway for positive regulation of gene transcription from a receptor on the cell surface (Soares-Silva et al., 2016). The presence of a PPP Fe-Zn-dependent phosphatase (FC 1.8) involved in reversible protein posttranslational modification points also in this direction. Cytosolic Hsp70 chaperone (FC 12.422) may also be involved in this MAPK signaling pathway, as is the case in mammals (Fan et al., 2018). GTPase activating component Ran-GAP (FC 2.862) is also known to be implicated downstream of MAPK responses (Faustino et al., 2007). A number of proteins involved in different stages of synthesis and maturation of RNAs and proteins are found. This includes the mitochondrial Tr-type G domain-containing GTPase/elongation factor Tu (FC 8.245) which bring the aminoacyl-tRNA into the A site of the mitoribosome, and Nsa1/WDR74 (FC 3.966), an assembly factor involved in the maturation of the large subunit of the cytosolic ribosome. The only ribosomal protein that is overaccumulated is RPL38, suggesting that it performs an additional function. Several proteins involved in the biogenesis and maturation of mRNA and ribosomes are found at FC values of 2-3, which are at the ‘executive’ side of signal transduction pathways that end in protein synthesis (Table I).

Of note are the three plastidial small heat shock proteins (HSP20, FC 12.510, 10.588, 6.530) for which little data exist in microalgae but in plants play a central role in the protection against stress damage, in the folding, intracellular distribution, and degradation of proteins, as well as in signal transduction chains (Ouyang et al., 2009). The HSPs are known to be generally involved in the response to stress most notably due to heat, but also other stresses that can affect protein stability such as oxidative stress, salinity or pH (Strauch and Haslbeck, 2016). Acetate is more likely than butyrate to be transported across the cell and imported into organelles to give rise to acetyl-CoA and further biosynthesis reactions (Boyle et al., 2017). Butyrate is likely only taken up in the peroxisome and di/tricarboxylates are exported into the cytoplasm and further into organelles. Since acetate is transported in the protonated form it will systematically release H+ within the organelles, which may cause some level of stress and possibly explain the increased need for HSP20.

The mitochondrial organization/maturation factor (CHCH domain) (FC 16.131) is the most induced protein compared to butyrate. Its function is uncertain, but may relate to protein translation or post-translational maturation of cytochrome c oxidase. Finally, although most enzymes involved in amino acid synthesis were mildly induced by butyrate, a few enzymes involved in production of aromatic amino acids and methionine were more abundant on acetate (FC ∼2.5). Methionine is a direct precursor of S-adenosylmethionine (SAM), an important posttranslational regulator of many cellular processes, including autophagy, the recycling of cellular components in response to stress (Ouyang et al., 2020). Butyrate induction of proteins such as the stress-induced carboxypeptidase (SCPL, FC 0.47) (Xu et al., 2021) and universal stress protein (IMP2, FC 0.34) seem to indicate that butyrate indeed causes some level of stress to the cells. It may thus be hypothesized that butyrate causes a decrease methionine to lower SAM, since it inhibits processes involved in stress response such as autophagy.

## 4 Conclusions and perspectives

The key hypotheses that have been proposed in the past concerning limitations in the trophic metabolism of microalgae relate to cell permeability, toxic products formed from the substrate, lack of enzymes necessary for effective dissimilation of the substrate or their improper cellular location, lack of transcriptional control, effect of low-intensity light in stimulating heterotrophic growth and respiratory deficiency, etc. (Neilson and Lewin, 1974). In this work, these key hypotheses were considered with regard to butyrate assimilation in *Polytomella* sp., and the relation with butyrate metabolism in other organisms is discussed as well as the potential implications of these findings for the capacities for butyrate assimilation of other -green-algae. In addition, a major step has been made in our understanding of peroxisomes in *Polytomella* sp. and in relation to its close relative *C. reinhardtii*.

Based on our data, we propose that butyrate is assimilated via peroxisomal β-oxidation resulting in acetyl-CoA and di/tricarboxylates for cellular use via the glyoxylate cycle. We found that multiple transporters are induced to facilitate the metabolic interplay between peroxisome and other cell compartments. Although no monocarboxylate transporter was identified for butyrate transport, a formate/nitrite transporter was put forward as candidate for this function. We hypothesize that butyrate causes a major antioxidant defense response related to the production of H_2_O_2_ and NADH in β-oxidation. An increased turnover of BCAAs to propionyl-CoA and acetyl-CoA was suggested, which may, together with an overproduction of pyruvate from glycolysis, serve amino acid or cofactor production. Butyrate lowers accumulation of carbohydrates and lipids while fatty acid synthesis was found induced, probably in the mitochondria. This all may serve organellar reorganization (peroxisomes) and the production of cofactors for several central metabolic enzymes to accommodate butyrate utilization. In contrast, acetate utilization seems to stimulate activities that relate to the biosynthesis and homeostasis of proteins.

Its fast butyrate assimilation makes *Polytomella* sp. a good model for the study of VFA metabolism, but the high starch levels even during the exponential growth is another distinctive trait that allow continuous cultivation on dark fermentation effluents with potential for biofuel production. Our proteomics approach revealed in many instances the induction on butyrate of multiple proteins belonging to the same pathway or similar metabolic activities suggest their importance in butyrate metabolism. Other omics and biochemical approaches should be employed to further explore the specificities of butyrate vs acetate as a carbon source. In particular, metabolomics and fluxomics should be used to reveal the assimilation pathways. The main issues that remain to be tackled relate to the import of VFAs into the cell and the role of β-oxidation and associated antioxidant activities, especially in green algae. As a non-photosynthetic alga, *Polytomella* can serve as a metabolic reference for efficient butyrate assimilation, to which other (green) algae may be compared. Since *Polytomella* sp. does not seem to appear to possess novel metabolic capacities *per se*, it should be considered that this alga owes its fast butyrate assimilation in some way to the loss of another major metabolic capacity: photosynthesis.

## 7 Author Contributions

JL performed experiments, data acquisition, data curation, formal analysis and writing of the original draft. AA contributed to conceptualization, experiments, data acquisition, data curation, formal analysis, validation, review and editing of the original draft. SB performed experiments, data acquisition, data curation and formal analysis. YC performed data acquisition, data curation, formal analysis, validation, review and editing of the original draft. OV performed data acquisition, data curation, formal analysis, validation, review and editing of the original draft. JPS contributed to supervision, funding acquisition, validation, review and editing of the original draft. RVL designed original experimental plan and performed experiments, data acquisition, data curation, formal analysis, supervision, funding acquisition, validation, review and editing of the original draft. All authors approved the final version of the manuscript.

## 10 Supplementary Material

Supplementary Table I. Differential analysis of total proteomes from *Polytomella* sp. grown on acetate or butyrate.

## 11 Data Availability Statement

The mass spectrometry proteomics data have been deposited to the ProteomeXchange Consortium via the PRIDE partner repository with the dataset identifier PXD035155 (https://www.ebi.ac.uk/pride/login)

## References

Acién Fernández, F. G., Fernández Sevilla, J. M., and Molina Grima, E. (2019). “Costs analysis of microalgae production,” in Biofuels from Algae (Elsevier), 551–566. doi: 10.1016/B978-0-444-64192-2.00021-4.

Adams, C., Godfrey, V., Wahlen, B., Seefeldt, L., and Bugbee, B. (2013). Understanding precision nitrogen stress to optimize the growth and lipid content tradeoff in oleaginous green microalgae. Bioresour. Technol. 131, 188–194. doi: 10.1016/j.biortech.2012.12.143.

Alban, C., Job, D., and Douce, R. (2000). BIOTIN METABOLISM IN PLANTS. Annu. Rev. Plant Physiol. Plant Mol. Biol. 51, 17–47. doi: 10.1146/annurev.arplant.51.1.17.

Almagro Armenteros, J. J., Sønderby, C. K., Sønderby, S. K., Nielsen, H., and Winther, O. (2017). DeepLoc: prediction of protein subcellular localization using deep learning. Bioinformatics 33, 3387–3395. doi: 10.1093/bioinformatics/btx431.

Arent, S., Christensen, C. E., Pye, V. E., Nørgaard, A., and Henriksen, A. (2010). The Multifunctional Protein in Peroxisomal β-Oxidation. J. Biol. Chem. 285, 24066–24077. doi: 10.1074/jbc.M110.106005.

Binder, S. (2010). Branched-Chain Amino Acid Metabolism in Arabidopsis thaliana. Arab. B. 8, e0137. doi: 10.1199/tab.0137.

Bouyssié, D., Hesse, A.-M., Mouton-Barbosa, E., Rompais, M., Macron, C., Carapito, C., et al. (2020). Proline: an efficient and user-friendly software suite for large-scale proteomics. Bioinformatics. doi: 10.1093/bioinformatics/btaa118.

Boyle, N. R., Sengupta, N., and Morgan, J. A. (2017). Metabolic flux analysis of heterotrophic growth in Chlamydomonas reinhardtii. PLoS One 12, 1–23. doi: 10.1371/journal.pone.0177292.

Bricker, D. K., Taylor, E. B., Schell, J. C., Orsak, T., Boutron, A., Chen, Y.-C., et al. (2012). A Mitochondrial Pyruvate Carrier Required for Pyruvate Uptake in Yeast, Drosophila, and Humans. Science (80-.). 337, 96–100. doi: 10.1126/science.1218099.

Casabona, M. G., Vandenbrouck, Y., Attree, I., and Couté, Y. (2013). Proteomic characterization of Pseudomonas aeruginosa PAO1 inner membrane. Proteomics 13, 2419–23. doi: 10.1002/pmic.201200565.

Casal, M., and Leão, C. (1995). Utilization of short-chain monocarboxylic acids by the yeast Torulaspora delbrueckii: Specificity of the transport systems and their regulation. BBA - Mol. Cell Res. 1267, 122–130. doi: 10.1016/0167-4889(95)00067-3.

Caverzan, A., Passaia, G., Rosa, S. B., Ribeiro, C. W., Lazzarotto, F., and Margis-Pinheiro, M. (2012). Plant responses to stresses: Role of ascorbate peroxidase in the antioxidant protection. Genet. Mol. Biol. 35, 1011–1019. doi: 10.1590/S1415-47572012000600016.

Chalima, A., Oliver, L., Fernández de Castro, L., Karnaouri, A., Dietrich, T., and Topakas, E. (2017). Utilization of Volatile Fatty Acids from Microalgae for the Production of High Added Value Compounds. Fermentation 3, 54. doi: 10.3390/fermentation3040054.

Choi, H. Il, Hwang, S. W., Kim, J., Park, B., Jin, E. S., Choi, I. G., et al. (2021). Augmented CO2 tolerance by expressing a single H+-pump enables microalgal valorization of industrial flue gas. Nat. Commun. 12, 1–16. doi: 10.1038/s41467-021-26325-5.

Craig, R. J., Hasan, A. R., Ness, R. W., and Keightley, P. D. (2021). Comparative genomics of Chlamydomonas. Plant Cell 33, 1016–1041. doi: 10.1093/plcell/koab026.

Cuff, M., Dyer, J., Jones, M., and Shirazi-Beechey, S. (2005). The human colonic monocarboxylate transporter Isoform 1: Its potential importance to colonic tissue homeostasis. Gastroenterology 128, 676–686. doi: 10.1053/j.gastro.2004.12.003.

Dakshinamurti, K. (2005). Biotin - A regulator of gene expression. J. Nutr. Biochem. 16, 419–423. doi: 10.1016/j.jnutbio.2005.03.015.

Davidi, L., Levin, Y., Ben-Dor, S., and Pick, U. (2014). Proteome Analysis of Cytoplasmatic and Plastidic β-Carotene Lipid Droplets in Dunaliella bardawil. Plant Physiol. 167, 60–79. doi: 10.1104/pp.114.248450.

De Duve, C., and Baudhuin, P. (1966). Peroxisomes (microbodies and related particles). Physiol. Rev. 46, 323–357. doi: 10.1152/physrev.1966.46.2.323.

de la Cruz, V. F., and Gittleson, S. M. (1981). The genus Polytomella: A review of classification, morphology, life cycle, metabolism, and motility. Arch. fur Protistenkd. 124, 1–28. doi: 10.1016/S0003-9365(81)80001-2.

De Meur, Q., Deutschbauer, A., Koch, M., Bayon-Vicente, G., Cabecas Segura, P., Wattiez, R., et al. (2020). New perspectives on butyrate assimilation in Rhodospirillum rubrum S1H under photoheterotrophic conditions. BMC Microbiol. 20, 1–20. doi: 10.1186/s12866-020-01814-7.

De Preter, V., Arijs, I., Windey, K., Vanhove, W., Vermeire, S., Schuit, F., et al. (2012). Impaired butyrate oxidation in ulcerative colitis is due to decreased butyrate uptake and a defect in the oxidation pathway*. Inflamm. Bowel Dis. 18, 1127–1136. doi: 10.1002/ibd.21894.

Delrue, F., Álvarez-Díaz, P., Fon-Sing, S., Fleury, G., and Sassi, J.-F. (2016). The Environmental Biorefinery: Using Microalgae to Remediate Wastewater, a Win-Win Paradigm. Energies 9, 132. doi: 10.3390/en9030132.

Dincer, I., and Acar, C. (2015). Review and evaluation of hydrogen production methods for better sustainability. Int. J. Hydrogen Energy 40, 11094–11111. doi: 10.1016/j.ijhydene.2014.12.035.

Donohoe, D. R., Collins, L. B., Wali, A., Bigler, R., Sun, W., and Bultman, S. J. (2012). The Warburg Effect Dictates the Mechanism of Butyrate-Mediated Histone Acetylation and Cell Proliferation. Mol. Cell 48, 612–626. doi: 10.1016/j.molcel.2012.08.033.

Duncan, S. H., Barcenilla, A., Stewart, C. S., Pryde, S. E., and Flint, H. J. (2002). Acetate Utilization and Butyryl Coenzyme A (CoA):Acetate-CoA Transferase in Butyrate-Producing Bacteria from the Human Large Intestine. Appl. Environ. Microbiol. 68, 5186–5190. doi: 10.1128/AEM.68.10.5186-5190.2002.

Durante, L., Hübner, W., Lauersen, K. J., and Remacle, C. (2019). Characterization of the GPR1/FUN34/YaaH protein family in the green microalga Chlamydomonas suggests their role as intracellular membrane acetate channels. Plant Direct 3. doi: 10.1002/pld3.148.

Fan, W., Gao, X. K., Rao, X. S., Shi, Y. P., Liu, X. C., Wang, F. Y., et al. (2018). Hsp70 Interacts with Mitogen-Activated Protein Kinase (MAPK)-Activated Protein Kinase 2 To Regulate p38MAPK Stability and Myoblast Differentiation during Skeletal Muscle Regeneration. Mol. Cell. Biol. 38. doi: 10.1128/MCB.00211-18.

Faustino, R. S., Stronger, L. N. W., Richard, M. N., Czubryt, M. P., Ford, D. A., Prociuk, M. A., et al. (2007). RanGAP-Mediated Nuclear Protein Import in Vascular Smooth Muscle Cells Is Augmented by Lysophosphatidylcholine. Mol. Pharmacol. 71, 438–445. doi: 10.1124/mol.105.021667.

Fei, Q., Fu, R., Shang, L., Brigham, C. J., and Chang, H. N. (2015). Lipid production by microalgae Chlorella protothecoides with volatile fatty acids (VFAs) as carbon sources in heterotrophic cultivation and its economic assessment. Bioprocess Biosyst. Eng. 38, 691–700. doi: 10.1007/s00449-014-1308-0.

Fleming, S. E., Fitch, M. D., DeVries, S., Liu, M. L., and Kight, C. (1991). Nutrient Utilization by Cells Isolated from Rat Jejunum, Cecum and Colon. J. Nutr. 121, 869–878. doi: 10.1093/jn/121.6.869.

Fuentes-Ramírez, E. O., Vázquez-Acevedo, M., Cabrera-Orefice, A., Guerrero-Castillo, S., and González-Halphen, D. (2021). The plastid proteome of the nonphotosynthetic chlorophycean alga Polytomella parva. Microbiol. Res. 243. doi: 10.1016/j.micres.2020.126649.

Gabaldón, T. (2010). Peroxisome diversity and evolution. Philos. Trans. R. Soc. B Biol. Sci. 365, 765–773. doi: 10.1098/rstb.2009.0240.

Garrison, R. G., Mirikitani, F. K., Henry, D. P., Evans, B. J., and Arnold, W. N. (1985). Ultrastructure of Candida ingens: a yeast that can assimilate volatile fatty acids. Microbios 42, 77–89.

Gerhardt, B. (1971). Zur Lokalisation von Enzymen der Microbodies in Polytomella caeca. Arch. Mikrobiol. 80, 205–218. doi: 10.1007/BF00410122.

Gerhardt, B., and Berger, C. (1971). Microbodies und Diaminobenzidin-Reaktion in den Acetat-Flagellaten Polytomella caeca und Chlorogonium elongatum. Planta 100, 155–166. Available at: http://sagdb.uni-goettingen.de/showstrains.php?allfields=caeca&strain_number=&prev_name=&genus=&division=&species=&clas=&search=Show+strains1.

Gonzalez, N. H., Felsner, G., Schramm, F. D., Klingl, A., Maier, U.-G., and Bolte, K. (2011). A Single Peroxisomal Targeting Signal Mediates Matrix Protein Import in Diatoms. PLoS One 6, e25316. doi: 10.1371/journal.pone.0025316.

Guan, X., Okazaki, Y., Zhang, R., Saito, K., and Nikolaua, B. J. (2020). Dual-localized enzymatic components constitute the fatty acid synthase systems in mitochondria and plastids. Plant Physiol. 183, 517–529. doi: 10.1104/pp.19.01564.

Haigh, W. G., and Beevers, H. (1964). The Glyoxylate cycle in Polytomella caeca. Arch. Biochem. Biophys. 107, 152–157.

Harris, E. H. (1989). The Chlamydomonas sourcebook.

Harris, E. H. (2001). Chlamydomonas as a model organism. Annu. Rev. Plant Physiol. Plant Mol. Biol. 52, 363–406. doi: 10.1146/annurev.arplant.52.1.363.

Hettema, E. H., van Roermund, C. W., Distel, B., van den Berg, M., Vilela, C., Rodrigues-Pousada, C., et al. (1996). The ABC transporter proteins Pat1 and Pat2 are required for import of long-chain fatty acids into peroxisomes of Saccharomyces cerevisiae. EMBO J. 15, 3813–22. Available at: http://www.ncbi.nlm.nih.gov/pubmed/8670886.

Hosotani, K., Ohkochi, T., Inui, H., Yokota, A., Nakano, Y., and Kitaoka, S. (1988). Photoassimilation of Fatty Acids, Fatty Alcohols and Sugars by Euglena gracilis Z. Microbiology 134, 61–66. doi: 10.1099/00221287-134-1-61.

Hu, J., Baker, A., Bartel, B., Linka, N., Mullen, R. T., Reumann, S., et al. (2012). Plant Peroxisomes: Biogenesis and Function. Plant Cell 24, 2279–2303. doi: 10.1105/tpc.112.096586.

Huerta-Cepas, J., Szklarczyk, D., Heller, D., Hernández-Plaza, A., Forslund, S. K., Cook, H., et al. (2019). eggNOG 5.0: a hierarchical, functionally and phylogenetically annotated orthology resource based on 5090 organisms and 2502 viruses. Nucleic Acids Res. 47, D309–D314. doi: 10.1093/nar/gky1085.

Hutner, S. H. (1972). Inorganic Nutrition. Annu. Rev. Microbiol. 26, 313–346. doi: 10.1146/annurev.mi.26.100172.001525.

Janssen, P. H., and Schink, B. (1995). Pathway of butyrate catabolism by Desulfobacterium cetonicum. J. Bacteriol. 177, 3870–3872. doi: 10.1128/jb.177.13.3870-3872.1995.

Johnson, X., and Alric, J. (2013). Central carbon metabolism and electron transport in chlamydomonas reinhardtii: Metabolic constraints for carbon partitioning between oil and starch. Eukaryot. Cell 12, 776–793. doi: 10.1128/EC.00318-12.

Karnaouri, A., Chalima, A., Kalogiannis, K. G., Varamogianni-Mamatsi, D., Lappas, A., and Topakas, E. (2020). Utilization of lignocellulosic biomass towards the production of omega-3 fatty acids by the heterotrophic marine microalga Crypthecodinium cohnii. Bioresour. Technol. 303, 122899. doi: 10.1016/j.biortech.2020.122899.

Kato, N., Nelson, G., and Lauersen, K. J. (2021). Subcellular Localizations of Catalase and Exogenously Added Fatty Acid in Chlamydomonas reinhardtii. Cells 10, 1940. Available at: https://www.mdpi.com/2073-4409/10/8/1940.

Kong, F., Liang, Y., Légeret, B., Beyly-Adriano, A., Blangy, S., Haslam, R. P., et al. (2017). Chlamydomonas carries out fatty acid β-oxidation in ancestral peroxisomes using a bona fide acyl-CoA oxidase. Plant J. 90, 358–371. doi: 10.1111/tpj.13498.

Krieger-Liszkay, A., and Feilke, K. (2016). The Dual Role of the Plastid Terminal Oxidase PTOX: Between a Protective and a Pro-oxidant Function. Front. Plant Sci. 6. doi: 10.3389/fpls.2015.01147.

Kunze, M., Pracharoenwattana, I., Smith, S. M., and Hartig, A. (2006). A central role for the peroxisomal membrane in glyoxylate cycle function. Biochim. Biophys. Acta - Mol. Cell Res. 1763, 1441–1452. doi: 10.1016/j.bbamcr.2006.09.009.

Lacroux, J., Seira, J., Trably, E., Bernet, N., Steyer, J., and van Lis, R. (2021). Mixotrophic Growth of Chlorella sorokiniana on Acetate and Butyrate : Interplay Between Substrate, C : N Ratio and pH. Front. Microbiol. 12, 1–16. doi: 10.3389/fmicb.2021.703614.

Lacroux, J., Trably, E., Bernet, N., Steyer, J. P., and van Lis, R. (2020). Mixotrophic growth of microalgae on volatile fatty acids is determined by their undissociated form. Algal Res. 47, 101870. doi: 10.1016/j.algal.2020.101870.

Lauersen, K. J., Willamme, R., Coosemans, N., Joris, M., Kruse, O., and Remacle, C. (2016). Peroxisomal microbodies are at the crossroads of acetate assimilation in the green microalga Chlamydomonas reinhardtii. Algal Res. 16, 266–274. doi: 10.1016/j.algal.2016.03.026.

Li-Beisson, Y., Beisson, F., and Riekhof, W. (2015). Metabolism of acyl-lipids in Chlamydomonas reinhardtii. Plant J. 82, 504–522. doi: 10.1111/tpj.12787.

Li-Beisson, Y., Shorrosh, B., Beisson, F., Andersson, M. X., Arondel, V., Bates, P. D., et al. (2013). Acyl-Lipid Metabolism. Arab. B. 11, e0161. doi: 10.1199/tab.0161.

Li-Beisson, Y., Thelen, J. J., Fedosejevs, E., and Harwood, J. L. (2019). The lipid biochemistry of eukaryotic algae. Prog. Lipid Res. 74, 31–68. doi: 10.1016/j.plipres.2019.01.003.

Liang, Y., Kong, F., Torres-Romero, I., Burlacot, A., Cuine, S., Légeret, B., et al. (2019). Branched-Chain Amino Acid Catabolism Impacts Triacylglycerol Homeostasis in Chlamydomonas reinhardtii. Plant Physiol. 179, 1502–1514. doi: 10.1104/pp.18.01584.

Liebthal, M., Maynard, D., and Dietz, K.-J. (2018). Peroxiredoxins and Redox Signaling in Plants. Antioxid. Redox Signal. 28, 609–624. doi: 10.1089/ars.2017.7164.

Linka, N., and Theodoulou, F. L. (2013). “Metabolite Transporters of the Plant Peroxisomal Membrane: Known and Unknown,” in, 169–194. doi: 10.1007/978-94-007-6889-5_10.

Liu, C. H., Chang, C. Y., Liao, Q., Zhu, X., and Chang, J. S. (2013). Photoheterotrophic growth of Chlorella vulgaris ESP6 on organic acids from dark hydrogen fermentation effluents. Bioresour. Technol. 145, 331–336. doi: 10.1016/j.biortech.2012.12.111.

Llamas, M., Dourou, M., González-Fernández, C., Aggelis, G., and Tomás-Pejó, E. (2020). Screening of oleaginous yeasts for lipid production using volatile fatty acids as substrate. Biomass and Bioenergy 138, 1–10. doi: 10.1016/j.biombioe.2020.105553.

Lloyd, D., Evans, D. A., and Venables, S. E. (1968). Propionate assimilation in the flagellate Polytomella caeca. An inducible mitochondrial enzyme system. Biochem. J. 109, 897–907. doi: 10.1042/bj1090897.

Lu, W., Du, J., Schwarzer, N. J., Gerbig-Smentek, E., Einsle, O., and Andrade, S. L. A. (2012). The formate channel FocA exports the products of mixed-acid fermentation. Proc. Natl. Acad. Sci. 109, 13254–13259. doi: 10.1073/pnas.1204201109.

May, P., Wienkoop, S., Kempa, S., Usadel, B., Christian, N., Rupprecht, J., et al. (2008). Metabolomics- and Proteomics-Assisted Genome Annotation and Analysis of the Draft Metabolic Network of Chlamydomonas reinhardtii. Genetics 179, 157–166. doi: 10.1534/genetics.108.088336.

McCommis, K. S., and Finck, B. N. (2015). Mitochondrial pyruvate transport: a historical perspective and future research directions. Biochem. J. 466, 443–454. doi: 10.1042/BJ20141171.

Mentzen, W. I., Peng, J., Ransom, N., Nikolau, B. J., and Wurtele, E. S. (2008). Articulation of three core metabolic processes in Arabidopsis: Fatty acid biosynthesis, leucine catabolism and starch metabolism. BMC Plant Biol. 8, 76. doi: 10.1186/1471-2229-8-76.

Millar, A. H., and Heazlewood, J. L. (2003). Genomic and Proteomic Analysis of Mitochondrial Carrier Proteins in Arabidopsis. Plant Physiol. 131, 443–453. doi: 10.1104/pp.009985.

Mishra, S. K., Suh, W. I., Farooq, W., Moon, M., Shrivastav, A., Park, M. S., et al. (2014). Rapid quantification of microalgal lipids in aqueous medium by a simple colorimetric method. Bioresour. Technol. 155, 330–333. doi: 10.1016/j.biortech.2013.12.077.

Morth, J. P., Pedersen, B. P., Buch-Pedersen, M. J., Andersen, J. P., Vilsen, B., Palmgren, M. G., et al. (2011). A structural overview of the plasma membrane Na+,K+-ATPase and H+-ATPase ion pumps. Nat. Rev. Mol. Cell Biol. 12, 60–70. doi: 10.1038/nrm3031.

Moscoviz, R., Trably, E., Bernet, N., and Carrère, H. (2018). The environmental biorefinery: State-of-the-art on the production of hydrogen and value-added biomolecules in mixed-culture fermentation. Green Chem. 20, 3159–3179. doi: 10.1039/c8gc00572a.

Neilson, A. H., and Lewin, R. A. (1974). The uptake and utilization of organic carbon by algae: an essay in comparative biochemistry. Phycologia 13, 227–264. doi: 10.2216/i0031-8884-13-3-227.1.

Nogaj, L. A., Srivastava, A., van Lis, R., and Beale, S. I. (2005). Cellular levels of glutamyl-tRNA reductase and glutamate-1-semialdehyde aminotransferase do not control chlorophyll synthesis in Chlamydomonas reinhardtii. Plant Physiol. 139, 389–396. doi: 10.1104/pp.105.067009.

Nolte, J. C., Schürmann, M., Schepers, C. L., Vogel, E., Wübbeler, J. H., and Steinbüchel, A. (2014). Novel characteristics of succinate coenzyme a (succinate-coa) ligases: Conversion of malate to malyl-coa and coa-thioester formation of succinate analogues in vitro. Appl. Environ. Microbiol. 80, 166–176. doi: 10.1128/AEM.03075-13.

Ouyang, Y., Chen, J., Xie, W., Wang, L., and Zhang, Q. (2009). Comprehensive sequence and expression profile analysis of Hsp20 gene family in rice. Plant Mol. Biol. 70, 341–357. doi: 10.1007/s11103-009-9477-y.

Ouyang, Y., Wu, Q., Li, J., Sun, S., and Sun, S. (2020). S-adenosylmethionine: A metabolite critical to the regulation of autophagy. Cell Prolif. 53. doi: 10.1111/cpr.12891.

Pan, R., Liu, J., Wang, S., and Hu, J. (2020). Peroxisomes: versatile organelles with diverse roles in plants. New Phytol. 225, 1410–1427. doi: 10.1111/nph.16134.

Pan, Y., Yang, J., Gong, Y., Li, X., and Hu, H. (2017). 3-Hydroxyisobutyryl-CoA hydrolase involved in isoleucine catabolism regulates triacylglycerol accumulation in Phaeodactylum tricornutum. Philos. Trans. R. Soc. B Biol. Sci. 372, 20160409. doi: 10.1098/rstb.2016.0409.

Perez-Garcia, O., and Bashan, Y. (2015). “Microalgal Heterotrophic and Mixotrophic Culturing for Bio-refining: From Metabolic Routes to Techno-economics,” in Algal Biorefineries (Cham: Springer International Publishing), 61–131. doi: 10.1007/978-3-319-20200-6_3.

Perez-Garcia, O., Escalante, F. M. E., de-Bashan, L. E., and Bashan, Y. (2011). Heterotrophic cultures of microalgae: Metabolism and potential products. Water Res. 45, 11–36. doi: 10.1016/J.WATRES.2010.08.037.

Perez-Riverol, Y., Bai, J., Bandla, C., García-Seisdedos, D., Hewapathirana, S., Kamatchinathan, S., et al. (2022). The PRIDE database resources in 2022: a hub for mass spectrometry-based proteomics evidences. Nucleic Acids Res. 50, D543–D552. doi: 10.1093/nar/gkab1038.

Petrova, V. Y., Drescher, D., Kujumdzieva, A. V., and Schmitt, M. J. (2004). Dual targeting of yeast catalase A to peroxisomes and mitochondria. Biochem. J. 380, 393–400. doi: 10.1042/bj20040042.

Pietrocola, F., Galluzzi, L., Bravo-San Pedro, J. M., Madeo, F., and Kroemer, G. (2015). Acetyl coenzyme A: a central metabolite and second messenger. Cell Metab. 21, 805–821. doi: 10.1016/j.cmet.2015.05.014.

Rezayian, M., Niknam, V., and Ebrahimzadeh, H. (2019). Oxidative damage and antioxidative system in algae. Toxicol. Reports 6, 1309–1313. doi: 10.1016/j.toxrep.2019.10.001.

Riekhof, W. R., Sears, B. B., and Benning, C. (2005). Annotation of genes involved in glycerolipid biosynthesis in Chlamydomonas reinhardtii: Discovery of the betaine lipid synthase BTA1Cr. Eukaryot. Cell 4, 242–252. doi: 10.1128/EC.4.2.242-252.2005.

Roediger, W. E. (1982). Utilization of nutrients by isolated epithelial cells of the rat colon. Gastroenterology 83, 424–429.

Rottensteiner, H., and Theodoulou, F. L. (2006). The ins and outs of peroxisomes: Co-ordination of membrane transport and peroxisomal metabolism. Biochim. Biophys. Acta - Mol. Cell Res. 1763, 1527–1540. doi: 10.1016/j.bbamcr.2006.08.012.

Round, F. (1980). The evolution of pigmented and unpigmented unicells - a reconsideration of the Protista. BioSystems 12, 61–69.

Roy, U. K., Nielsen, B. V., and Milledge, J. J. (2021). Antioxidant production in Dunaliella. Appl. Sci. 11, 1–24. doi: 10.3390/app11093959.

Schwacke, R., Ponce-Soto, G. Y., Krause, K., Bolger, A. M., Arsova, B., Hallab, A., et al. (2019). MapMan4: A Refined Protein Classification and Annotation Framework Applicable to Multi-Omics Data Analysis. Mol. Plant 12, 879–892. doi: 10.1016/j.molp.2019.01.003.

Schwanhäusser, B., Busse, D., Li, N., Dittmar, G., Schuchhardt, J., Wolf, J., et al. (2011). Global quantification of mammalian gene expression control. Nature 473, 337–342. doi: 10.1038/nature10098.

Sheeler, P., Moore, J., Cantor, M., and Granik, R. (1968). The stored polysaccharide of Polytomella agilis. Life Sci. 7, 1045–1051. doi: 10.1016/0024-3205(68)90141-0.

Shtaida, N., Khozin-Goldberg, I., and Boussiba, S. (2015). The role of pyruvate hub enzymes in supplying carbon precursors for fatty acid synthesis in photosynthetic microalgae. Photosynth. Res. 125, 407–422. doi: 10.1007/s11120-015-0136-7.

Silverberg, B. A. (1975). An ultrastructural and cytochemical characterization of microbodies in the green algae. Protoplasma 83, 269–295. doi: 10.1007/BF01282559.

Smith, D. R., and Lee, R. W. (2014). A plastid without a genome: evidence from the nonphotosynthetic green algal genus Polytomella. Plant Physiol. 164, 1812–1819. doi: 10.1104/pp.113.233718.

Soares-Silva, M., Diniz, F. F., Gomes, G. N., and Bahia, D. (2016). The Mitogen-Activated Protein Kinase (MAPK) Pathway: Role in Immune Evasion by Trypanosomatids. Front. Microbiol. 7. doi: 10.3389/fmicb.2016.00183.

Sousa, R. H. V, Carvalho, F. E. L., Lima-Melo, Y., Alencar, V. T. C. B., Daloso, D. M., Margis-Pinheiro, M., et al. (2019). Impairment of peroxisomal APX and CAT activities increases protection of photosynthesis under oxidative stress. J. Exp. Bot. 70, 627–639. doi: 10.1093/jxb/ery354.

Strauch, A., and Haslbeck, M. (2016). The function of small heat-shock proteins and their implication in proteostasis. Essays Biochem. 60, 163–172. doi: 10.1042/EBC20160010.

Tardif, M., Atteia, A., Specht, M., Cogne, G., Rolland, N., Brugière, S., et al. (2012). PredAlgo: A New Subcellular Localization Prediction Tool Dedicated to Green Algae. Mol. Biol. Evol. 29, 3625–3639. doi: 10.1093/molbev/mss178.

Turon, V., Baroukh, C., Trably, E., Latrille, E., Fouilland, E., and Steyer, J.-P. (2015). Use of fermentative metabolites for heterotrophic microalgae growth: Yields and kinetics. Bioresour. Technol. 175, 342–349. doi: 10.1016/J.BIORTECH.2014.10.114.

Turon, V., Trably, E., Fouilland, E., and Steyer, J.-P. (2016). Potentialities of dark fermentation effluents as substrates for microalgae growth: A review. Process Biochem. 51, 1843–1854. doi: 10.1016/J.PROCBIO.2016.03.018.

van Lis, R., and Atteia, A. (2004). Control of Mitochondrial Function via Photosynthetic Redox Signals. Photosynth. Res. 79, 133–148. doi: 10.1023/B:PRES.0000015409.14871.68.

van Lis, R., Couté, Y., Brugière, S., Tourasse, N. J., Laurent, B., Nitschke, W., et al. (2021). Phylogenetic and functional diversity of aldehyde-alcohol dehydrogenases in microalgae. Plant Mol. Biol. 105, 497–511. doi: 10.1007/s11103-020-01105-9.

van Roermund, C. W. T., Schroers, M. G., Wiese, J., Facchinelli, F., Kurz, S., Wilkinson, S., et al. (2016). The Peroxisomal NAD Carrier from Arabidopsis Imports NAD in Exchange with AMP. Plant Physiol. 171, 2127–2139. doi: 10.1104/pp.16.00540.

van Roermund, C. W. T., Tabak, H. F., van den Berg, M., Wanders, R. J. A., and Hettema, E. H. (2000). Pex11p Plays a Primary Role in Medium-Chain Fatty Acid Oxidation, a Process That Affects Peroxisome Number and Size in Saccharomyces cerevisiae. J. Cell Biol. 150, 489–498. doi: 10.1083/jcb.150.3.489.

Walter, K. A., Nair, R. V., Cary, J. W., Bennett, G. N., and Papoutsakis, E. T. (1993). Sequence and arrangement of two genes of the butyrate-synthesis pathway of Clostridium acetobutylicum ATCC 824. Gene 134, 107–111. doi: 10.1016/0378-1119(93)90182-3.

Wieczorek, S., Combes, F., Lazar, C., Giai Gianetto, Q., Gatto, L., Dorffer, A., et al. (2017). DAPAR & ProStaR: software to perform statistical analyses in quantitative discovery proteomics. Bioinformatics 33, 135–136. doi: 10.1093/bioinformatics/btw580.

Wise, D. L. (1955). Carbon Sources for Polytomella caeca. J. Protozool. 2, 156–158. doi: 10.1111/j.1550-7408.1955.tb02416.x.

Wise, D. L. (1959). Carbon Nutrition and Metabolism of Polytomella caeca. J. Protozool. 6, 19–23. doi: 10.1111/j.1550-7408.1959.tb03921.x.

Wise, D. L. (1968). Effects of Acetaldehyde on Growth and Biosynthesis in an Algal Flagellate Polytomella caeca. J. Protozool. 15, 528–531. doi: 10.1111/j.1550-7408.1968.tb02169.x.

Wu, T., Fu, Y., Shi, Y., Li, Y., Kou, Y., Mao, X., et al. (2020). Functional Characterization of Long Chain Acyl-CoA Synthetase Gene Family from the Oleaginous Alga Chromochloris zofingiensis. J. Agric. Food Chem. 68, 4473–4484. doi: 10.1021/acs.jafc.0c01284.

Xu, X., Zhang, L., Zhao, W., Fu, L., Han, Y., Wang, K., et al. (2021). Genome-wide analysis of the serine carboxypeptidase-like protein family in Triticum aestivum reveals TaSCPL184-6D is involved in abiotic stress response. BMC Genomics 22, 350. doi: 10.1186/s12864-021-07647-6.

Yemm, E. W., and Willis, A. J. (1954). The estimation of carbohydrates in plant extracts by anthrone.Biochem. J. 57, 508–14. Available at: http://www.ncbi.nlm.nih.gov/pubmed/13181867.

Zhan, J., Rong, J., and Wang, Q. (2017). Mixotrophic cultivation, a preferable microalgae cultivation mode for biomass/bioenergy production, and bioremediation, advances and prospect. Int. J. Hydrogen Energy 42, 8505–8517. doi: 10.1016/j.ijhydene.2016.12.021.

